# Breeding responses to environmental variation are age- and trait-dependent in female Nazca boobies

**DOI:** 10.1101/2021.02.23.432506

**Authors:** Emily M. Tompkins, David J. Anderson

**Affiliations:** Biology Department, Wake Forest University, Winston-Salem, USA

**Keywords:** Age by environment interaction, breeding date, clutch size, diet, egg volume, Galápagos, El Niño-Southern Oscillation, La Niña, male age, seabird, senescence

## Abstract

Age and environment are important determinants of reproductive parameters in long-lived organisms. These factors may interact to determine breeding responses to environmental change, yet few studies have examined the environmental-dependence of aging patterns across the entire lifespan. We do so, using a 20-year longitudinal dataset of reproductive phenotypes in long-lived female Nazca boobies (*Sula granti*), a monogamous seabird breeding in the eastern tropical Pacific. Young and old females may suffer from inexperience and senescence, respectively, and/or practice reproductive restraint. Breeding performance (for breeding participation, breeding date, clutch size, egg volume, and offspring production) was expected to be lower in these age classes, particularly under environmental challenge, in comparison with middle-aged breeders. Sea surface temperature anomalies (SSTA) represented interannual variation in the El Niño-Southern Oscillation (ENSO) and were one proxy for environmental quality (a population count of clutch initiations was a second). Although only females lay eggs, both sexes care for eggs and nestlings, and the male partner’s age, alone or in interaction with female age, was evaluated as a predictor of breeding performance. Middle-aged females performed better than young and old birds for all reproductive traits. Pairing with a young male delayed breeding (particularly for old females) and reduced clutch size, and pairing with an old male reduced offspring production. Challenging environments increased age effects on breeding probability and breeding date across young to middle ages and for offspring production across middle to old ages. However, important exceptions to the predicted patterns for clutch size and fledging success across young to middle ages suggested trade-offs between fitness components may complicate patterns of trait expression across the lifespan. Relationships between breeding participation, environment, and individual quality and/or experience in young females may also contribute to unexpected patterns for clutch size and fledging success, traits expressed only in breeders. Finally, independent of age, breeding responses of female Nazca boobies to the ENSO did not follow expectations derived from oceanic forcing of primary productivity. During El Niño-like conditions, egg-laying traits (clutch size, breeding date) improved but offspring production declined, while La Niña-like conditions were “poor” environments throughout the breeding cycle.

## INTRODUCTION

Long-lived, iteroparous organisms confront substantial variation in the quality of the breeding environment during a lifespan. For these organisms, life history theory predicts a flexible response, tracking the environment via shifts in allocation of reproductive effort (Stearns 1992, Erikstad et al. 1998). Age, too, influences the constraints organisms face and their optimal allocation strategy (Williams 1966, Schaffer 1974, McNamara et al. 2009). At the intersection of these two factors, breeding responses to environmental quality can depend on age (Sydeman et al. 1991, Laaksonen et al. 2002). Quantifying age-dependent responses to environmental variation improves our understanding of how optimal behaviors and strategies change across the lifespan (McNamara et al. 2009) and is required for a mechanistic understanding of population fluctuations in long-lived species (e.g., Coulson et al. 2001), where age is a major factor structuring survival (reviewed in Nussey et al. 2013) and reproduction (reviewed in Lemaître and Gaillard 2017). Despite its relevance to understanding life history variation and population dynamics, studies evaluating age by environment interactions are still relatively rare, because, for many species, decades are required to observe old age classes in a range of environmental settings. Here, we use a multi-decade longitudinal dataset to evaluate links between reproductive phenotype, age, and interannual variation in the El Niño-Southern Oscillation (ENSO) in female Nazca boobies (*Sula granti*), a seabird breeding in the eastern tropical Pacific (ETP). We then explore interactive effects of age and environment on reproductive traits.

On the individual, longitudinal level, early-life improvement in performance could result from improved foraging or breeding competence with experience or growth (the “constraint” hypothesis; Nur 1984, Catry and Furness 1999), and/or low motivation in early life to risk allocation to current breeding when residual reproductive value is highest (the “restraint” hypothesis; Williams 1966, Curio 1983). These within-individual explanations for the relatively poor breeding performance of young individuals – inexperience or optimization of reproductive effort – each predict that young individuals perform particularly poorly in poor environments.

Therefore, age effects on breeding should be amplified under poor breeding environments and attenuated in good conditions. Accordingly, the pregnancy rates of yearling versus adult red deer (*Cervus elaphus*) diverge at high population density (Bonenfant et al. 2002) and the largest performance discrepancies between the offspring production of young versus middle-aged birds occur under low food availability in Western gulls (*Larus occidentalis*; Sydeman et al. 1991), and Australasian gannets (*Morus serrator*; Bunce et al. 2005). However, not all studies find age-dependent responses to environmental variation (e.g., Nevoux et al. 2007, Vieyra et al. 2009) and two studies have revealed the opposite pattern: poor environments shrink, not accentuate, age-related differences in breeding performance for Audouin’s gulls (*Larus audouinii*; Oro et al. 2014) and Scopoli’s shearwater (*Calonectris diomedea*; Hernández et al. 2015). Detecting within-individual aging patterns (the focus of this study), and evaluating their interaction with the environment, is best accomplished using longitudinal approaches that control individual heterogeneity (Cam et al. 2002, van de Pol and Verhulst 2006). Otherwise, processes acting among individuals (e.g., the selective disappearance of low-quality individuals from an aging cohort) may confound patterns of aging at the population level (Cam et al. 2002).

Turning to old age, reproductive senescence is common in mammals and birds (reviewed in Lemaître and Gaillard 2017), including seabirds (e.g., Kim et al. 2011, Froy et al. 2013), but is not ubiquitous (e.g., Sydeman et al. 1991, Berman et al. 2009). Age-related declines in reproductive parameters may reflect physiological senescence (documented in aging seabirds: Angelier et al. 2007, Elliott et al. 2014, but see Lecomte et al. 2010) and/or terminal restraint (McNamara et al. 2009). Terminal restraint is expected when physiological state, not age, sets an individual’s remaining lifespan. Under this hypothesis, reproductive activity in a poor environment advances biological age more than it would under good conditions and should be avoided (McNamara et al. 2009). A competing idea posits that old individuals, faced with dwindling opportunities for future reproduction, should make terminal investments (Clutton-Brock 1984). Both terminal investment (e.g., Velando et al. 2006, Froy et al. 2013) and terminal restraint (e.g., Elliott et al. 2014) have received empirical support. However, in Nazca boobies, any potential increase in reproductive investment in old age is apparently overwhelmed by senescence: old females perform much worse than middle-aged females, including for breeding probability (Tompkins and Anderson 2019). Thus, old Nazca boobies – constrained by physiological senescence or practicing adaptive restraint – are expected to show relatively greater reductions in performance, compared to middle-aged individuals, in poor environments.

We examined environment-specific age effects on reproductive performance of Nazca boobies with respect to the ENSO, a globally important climate phenomenon (Fiedler 2002; McPhaden et al. 2006). Nazca boobies forage for pelagic fish and squid (Anderson 1989, Tompkins et al. 2017) 10s-100s of km offshore in the ETP (Anderson and Ricklefs 1987, Zavalaga et al. 2012). This foraging area is in the equatorial “cold tongue”, a protrusion of usually cold and productive waters stretching west along the equator from the coast of South America that changes dramatically according to ENSO state. During an El Niño event, easterly trade winds slacken, and warm waters from the western Pacific move eastward, raising sea surface temperatures (SST) in the ETP and deepening the thermocline (a steep vertical gradient of SST; Wang and Fiedler 2006). Physical changes during El Niño reduce primary productivity in the ETP (Feldman et al. 1984, Pennington et al. 2006) and propagate through biological communities, influencing predators primarily through effects on prey species abundance or availability (Barber and Chavez 1983, Jahncke and Goya 2000, Ancona et al. 2012). Effects of extreme El Niño events on seabirds are typically negative: populations exhibit delayed breeding (Ancona et al. 2011) and reduced recruitment (Oro et al. 2010), breeding participation (Cubaynes et al. 2011), fledging success (Schreiber and Schreiber 1984, Anderson 1989, Ancona et al. 2011), juvenile survival (Champagnon et al. 2018), and/or adult survival (Oro et al. 2010). At the other ENSO extreme, easterly trade winds intensify during “La Niña” events, cooling eastern equatorial Pacific surface water and reducing thermocline depth. Some studies report enhanced primary productivity (Pennington et al. 2006) or reproductive success of seabirds when surface waters are cool (Oro et al. 2010, Ancona et al. 2011). However, important exceptions to predicted patterns (El Niño bad, La Niña good; e.g., Spear et al. 2001, 2007, Doherty et al. 2004, Devney et al. 2009), including in Nazca boobies (Champagnon et al. 2018), emphasize the need for a deeper understanding of the connections between trophic relationships, life histories, and breeding responses to oceanographic changes.

Causal relationships between ENSO phase and primary productivity imply that El Niño warm events should provide poor environments for breeding and La Niña cool events good ones. Surprisingly, the latter stage of breeding (rearing larger nestlings) and juvenile survival are affected negatively by both ENSO extremes in Nazca boobies (Champagnon et al. 2018, Tompkins and Anderson 2019), motivating investigation of ENSO effects during egg laying, which falls during the peak in ENSO-associated sea surface temperature anomalies (SSTA) in the ETP, both positive and negative. Early breeding and laying a second “insurance” egg (vs. a one-egg clutch) influence breeding success in Nazca boobies (Anderson 1990, Clifford and Anderson 2001) and trait expression is tightly linked to food availability, especially for clutch size (Clifford and Anderson 2001, 2002).

Our goal was to understand age-related performance across environmental variation, first by modelling egg-laying traits (breeding date, clutch size, and egg volume) and fledging success as a function of female age and of her male mate’s age (egg-laying traits are expressed in females but both sexes care for eggs and dependent young), and then by evaluating the extent to which female age interacts with environmental variation to shape breeding phenotypes in this booby population. Age- and environment-dependent breeding participation may influence patterns of variation in egg-laying traits and fledging success (traits expressed only in breeders). Age by environment interactions were also evaluated for female breeding probability to address this possibility. Environments associated with high average performance for each trait (early breeding and larger breeding probability, clutch size, egg volume, and fledging success) were expected to show smaller differences between young and middle-aged females, and between middle-aged and old females, compared to results from challenging environments. Few studies have examined how age interacts with oceanographic parameters to affect reproduction in seabirds; even fewer involve old age classes (Vieyra et al. 2009, Pardo et al. 2013, Oro et al. 2014). We fill an important empirical gap, and do so in the context of the ENSO, a climate phenomenon of global importance forecast to undergo changes in the character and frequency of extreme El Niño/La Niña phases under global climate change (Yeh et al. 2009, Cai et al. 2014).

## METHODS

### Data collection

Data on breeding probability, offspring production and egg-laying phenotypes (breeding date, clutch size, and volume of the first laid “A” egg) were collected from known-age female Nazca boobies at the Punta Cevallos colony on Isla Española in the Galápagos Islands (1°20’ S, 89°40’ W) from 1992-2005 and from 2010-2018. Nazca boobies at Punta Cevallos breed from October until May or June of the following calendar year. Breeding seasons are referenced by the first year in each two-year span and dates are referenced by extended Julian date, the number of days since Jan 1^st^ of the first calendar year. Since 1984, alphanumeric metal bands have identified individuals in the monitored area (Huyvaert and Anderson 2004) and allowed the accumulation of detailed longitudinal data on clutch initiations and breeding performance. Bands are read by capturing birds by hand at the nest site or using binoculars from a few meters distance.

All nests initiated by banded individuals within the monitored area (Huyvaert and Anderson 2004) were visited every other day (1992-1993 seasons) or daily (1994-2004, 2010-2017) from early in the breeding season through mid-January, when egg laying is effectively complete. “Breeding Probability” was defined as the probability of breeding, given that a focal female was alive, based on a female’s breeder/non-breeder status (from comprehensive nest monitoring) and alive/dead status (assessed from an annual band resight survey, details in Huyvaert and Anderson 2004). Recapture probabilities are > 0.85 for most ages/breeding seasons (Tompkins et al. 2017) and a female was considered “dead” in the first year of an absence of ≥ 2 consecutive years from the annual band resight survey (following Tompkins and Anderson 2019). Only 1% of banded birds reappear after a 2-year absence, requiring retrospective adjustment of year of death. “Breeding Date” was defined as the laying date of the first-laid egg in a clutch. “A-egg Volume” (in cm^3^) was calculated from measurements of egg length and breadth as 0.00051 * length * breadth^2^ + 1.22, following Clifford and Anderson (2002). “Clutch Size” is one or two eggs and was scored as a dichotomous trait and analysed as the probability of laying two eggs, given that a clutch was initiated. Egg-laying traits refer to the first-laid clutch in a season (a re-nest clutch is sometimes laid after a first clutch’s failure).

When two eggs hatch, two-nestling broods are reduced to one by obligate siblicide and “Fledging Success” was defined as the probability of raising a single offspring to independence, given that a nest was initiated. Fledging Success was based on a given female’s breeder/non-breeder status (from nest monitoring during egg laying) and survival of her offspring (banded in March-May) until independence. Offspring survival until independence is assessed through daily nest monitoring late in the breeding period (from March to late April or early May) and, because some offspring will not reach independence until June/July, is adjusted for any mortality occurring after monitoring ends but before offspring fledge (assessed by searching for banded offspring carcasses at the start of the following breeding season; see Appendix S1 for details and justification). Offspring from re-nest and original clutches were not distinguished. Additional details of data selection and adjustments to handle missing information are described in Appendix S1. Data for the current study comprise the breeding records of known-age females (cohorts 1984 and later) in seasons with detailed data on breeding attempts in the monitored area: 1993-2004 and 2010-2017. The data for Breeding Probability and Breeding Date exclude seasons 1998 and 1999 and 2017 (Breeding Probability only) because of an unusually high degree of missing information (Appendix S1).

The 20 years analyzed in this study include multiple El Niño and La Niña events (following the definition of Trenberth 1997; time series shown in Appendix S2). Interannual variation in the ENSO was quantified as the average December-February SSTA from the Niño 3 region (5°S-5°N, 150°W-90°W), calculated from monthly SSTA values downloaded from the LDEO/IRI data library (https://iridl.ldeo.columbia.edu, accessed April 19, 2020). The Niño 3 region includes the Nazca booby foraging range while rearing nestlings (Zavalaga et al. 2012) and interannual variation in ENSO state is highest during December-February (Cai et al. 2014), explaining our choice of location and time period. Average December-February SSTA from the Niño 3 region is highly correlated with SSTA during early egg laying (September-November, r = 0.97, d.f. = 24, p << 0.001, seasons 1992-2017), and during late chick rearing (March-May; r = 0.69, d.f. = 24, p << 0.001); thus, our measure of interannual variation in the ENSO reflects conditions experienced across the entire breeding cycle.

### Statistical analyses

#### Additive effects of female age and environment on egg-laying traits

Statistical analyses were conducted in a short sequence for each breeding response variable. First, additive effects of Female Age (continuous age in years) and environmental variables on egg-laying traits were examined (these relationships are already described for Breeding Probability and Fledging Success; Tompkins and Anderson 2019). “Age + Environment” analyses established (1) age-related patterns of trait expression for egg-laying traits and (2) relationships between SSTA and average performance. The latter revealed how the ENSO aligned with the quality of the breeding environment, with high performance (large Clutch Size and A-egg Volume, early breeding) assumed to signal a relatively resource-rich environment. To further evaluate a relationship between ENSO and resource availability, we fit a binomial GLMM to regurgitation data from systematic diet samples (N ≥ 20 per month in Nov., Dec., and Jan.), describing the effect of SSTA and SSTA^2^ on the probability of regurgitating one or more food items (data from seasons 1992-1994, 1999-2004, 2011-2017 only; details in Appendix S3). Age + Environment data included records from known-age females aged 3-25 yrs, covering much of the post-recruitment lifespan (ages 2 and > 25 not included because of small sample sizes). Sample sizes were > 14,300 records and > 2,645 females for all traits (sample sizes listed in Appendix S4).

We used a model selection approach to evaluate the importance of female age as a predictor of each egg-laying trait. The performance of candidate models featuring different functional relationships between age and breeding performance was compared using either 10-fold cross validation (10-fold CV; Hooten and Hobbs 2015) or approximate Leave-One-Out cross validation using Pareto Smoothed Importance Sampling (PSIS-LOO; Vehtari et al. 2016). Each approach evaluates and compares candidate models on the basis of their ability to make out-of-sample predictions using within-sample model fits. During 10-fold CV, predictive accuracy is scored by refitting each candidate model 10 times, each time leaving out 1/10^th^ of the data. Each set of left-out data is then compared with model predictions based on the other 9/10^th^ of the data to generate the expected log pointwise predictive density (*el^p^^d 10-fold CV*), a measure of predictive performance. Exact cross validation is computationally demanding. At its extreme, each of *n* data points would be left out in turn and the model would be refit *n* times (exact leave-one-out cross validation). Pareto Smoothed Importance Sampling approximates leave-one-out cross validation scores (*el^p^^d PSIS-LOO*), providing a robust estimate of out-of-sample prediction accuracy without refitting the model. We used *el^p^^d PSIS-LOO* as a basis for model comparison whenever a critical diagnostic (k-hat, the estimated shape parameter for the generalized Pareto distribution) indicated *el^p^^d PSIS-LOO* estimates were reliable, and 10-fold CV otherwise.

The most general parameterization of age effects modelled the relationship between egg-laying traits and Female Age using non-parametric smoothers (thin plate regression splines; Wood 2006) in Generalized Additive Mixed Models (GAMMs). Other candidate models (GLMMs) fit age as a linear function, a quadratic function, or using one- or two-threshold functions. These simpler parameterizations of age effects quantify rates of change in trait values with age, providing a test of early-life improvement and senescence. The best-supported candidate model among the simpler parameterizations of age effects was retained in subsequent analyses (described below). Two-threshold functions allowed the linear slope of performance on age to vary across three periods defined by two threshold ages (T_1_, T_2_): an early period (ages ≤ T_1_, where T_1_ = 6, 7, 8, or 9 years old in alternative candidate models), a prime-age period, and a late-life period (ages > T_2_, where T_2_ = 13, 14, 15, 16, or 17 years old in alternative candidate models). Single-threshold functions allowed the slope of performance with age to differ between an early period (ages ≤ T_1_, where T_1_ = 6, 7, 8, or 9) and all later ages (details of threshold model parameterizations are in Appendix S5). Threshold ages were chosen based on aging patterns for Breeding Probability and Fledging Success in this species (Tompkins and Anderson 2019).

Comparing the performance of single threshold models with that of two-threshold models gave the strength of evidence for a late-life change in aging rates. Comparing the performance of all candidate models including age effects to that of a model omitting age (but otherwise identical) indicated the importance of age-related changes in trait values.

Alongside age effects, SSTA was parameterized as a quadratic function, allowing peak egg-laying performance to associate with ENSO-neutral conditions (as observed for Fledging Success and juvenile survival in Nazca boobies; Tompkins and Anderson 2019, Champagnon et al. 2018). A continuous measure of the breeding population size (“Nest Count”) was a second index of environmental conditions. We assumed a positive association between Nest Count and the quality of the breeding environment, but a negative association is also plausible under density-dependent processes. In practice, a proxy for foraging trip length in incubating females was not affected by Nest Count (5 years of data, Appendix S6) and associations between Nest Count and breeding performance were positive (Clutch Size, Fledging Success) or not supported statistically, suggesting that any signal of intra-specific competition carried by Nest Count is less important than that of resource availability. Other predictors were a dichotomous factor marking a qualitative and quantitative shift in diet (“Fish Phase”; Tompkins et al. 2017), Fledging Success in the previous season (“FS_(*t*-1)_”, for each female), and relative breeding date (“rBD”; standardized within year; for Clutch Size and A-egg Volume only) to account for potential seasonal declines in performance. These explanatory variables are described in Table 1. Female Identity and Breeding Season were fit as group-specific (“random”) intercepts to account for individual differences in performance and the non-independence of records measured within the same breeding season. Clutch Size (and Breeding Probability and Fledging Success, in later stages) were modelled using a binomial error structure and a logit link function; models for all other response variables assumed Gaussian errors. GAMMs and GLMMs were fit in a Bayesian Framework using program Stan (Carpenter et al. 2017) and the rstanarm package (v. 2.13.1, Goodrich et al. 2016) in R (v. 4.0.1, R Core Team 2020). Details of priors, posterior estimation, and model fit are included in Appendix S7.

**Table 1.** Descriptions of variables (other than female age) used in analyses of age and environmental effects on Nazca booby breeding traits.

| Predictor(s) | Variable definition | Notes on interpretation or role in analyses |
| --- | --- | --- |
| SSTA, SSTA<sup>2</sup> | Average Dec.-Feb. monthly SSTA from 5°S-5°N, 150°W-90°W. | Positive sea surface temperature anomalies (SSTA) are "El Niño-like" and negative SSTA are "La Niña-like". Parameterized as a quadratic function to allow peak performance under neutral conditions (Champagnon et al. 2018, Tompkins et al. 2019). |
| Nest Count | Annual count of clutch initiations (excluding re-nests) in a colony subsection. | A measure of the quality of the breeding environment. SSTA and Nest Count are uncorrelated across the 20 years of our study ($r = 0.18$, d.f. = 18, $p = 0.45$). |
| Fish Phase | Dichotomous factor, levels: Sardine Phase (< 1997) and Flying Fish Phase ($\geq 1997$) | Maps a decadal-scale change in the diet of Nazca boobies from high quality sardine prey to low quality flying fish prey (Tompkins et al. 2017). |
| FS<sub>(t-1)</sub> | Dichotomous factor describing each female's Fledging Success the previous season (0,1). | In our data, breeding seasons with negative SSTA follow seasons of strongly positive SSTA (and of relatively low Fledging Success; Tompkins et al. 2019). FS<sub>(t-1)</sub> was included to separate the current influence of negative SSTA from any lagged effect of last season's breeding effort. |
| Earlier traits: rBD, rAV, CS | Relative (standardized within year) breeding date (rBD) and A-egg volume (rAV); dichotomous clutch size (1 vs. 2 eggs; CS) | Included to isolate environmental and age effects on Fledging Success from those acting on earlier-expressed traits potentially correlated with Fledging Success. For the same reason, rBD was a predictor of Clutch Size and A-egg Volume. |
| ALA | Age at last appearance in annual Band Resight Surveys (BRS), for females presumed dead (absent 2 or more years from the BRS). | A linear increase in performance with increasing lifespan is evidence for the selective disappearance of low-quality individuals with respect to egg-laying phenotypes. |
| M Age | Categorical male age. Young: $\leq 9$ years; Middle-aged: 10-15 years; Old: $\geq 16$ years. | Male age categories are defined based on thresholds defining early-life improvement and senescent decline in annual breeding success (Tompkins and Anderson 2019). |

From the candidate models with the best-supported parameterization of female age, the significance of other fixed effects was evaluated using the span of the 95% Bayesian credible interval (BCI) on each parameter estimate. Higher order terms (quadratic terms) with 95% BCI on coefficient estimates spanning zero were removed sequentially and simplified models re-run to verify the significance of lower-order terms. In this and later analyses, continuous independent variables (e.g., Nest Count) were standardized before analysis (zero mean, unit s.d.), except for SSTA, which naturally fell close to this scale. Female age was also re-scaled (divided by 10) to aid model convergence. Dependent variables Breeding Date and A-egg Volume were standardized across all years (preserving mean differences between years).

#### Selective disappearance

Early mortality of poor-quality individuals (e.g., Cam et al. 2002, Barbraud and Weimerskirch 2005) can result in the progressive improvement of average phenotypic quality, and thus performance, in an aging cohort. This “selective disappearance” can obscure or enhance age-related changes due to within-individual processes (e.g., learning, senescence). We evaluated the potential for selective disappearance to act in female Nazca boobies by estimating the effect of Age at Last Appearance (ALA, a proxy for lifespan) on Breeding Date, A-egg Volume, and Clutch Size (following van de Pol and Verhulst 2006; this effect is already established for Breeding Probability and Fledging Success by Tompkins and Anderson 2019). A positive correlation between age at last appearance and breeding performance suggests that poor quality individuals die young (selective disappearance is acting). The best-performing GLMM from the Age + Environment analyses (including Female Identity and Breeding Season random intercepts, as above) was modified to include ALA and fit to breeding records from known-age females belonging to cohorts 1984-1994. The data were restricted to these cohorts because 97.5% of the members are presumed dead (absent ≥ 2 years from annual band resight surveys). Ages at disappearance > 23 were assigned to 23, the minimum age of death for the 2.5% of females still “living”. Sample size varied by response variable (Appendix S4) but included > 3,000 cases from ≥ 470 unique females.

#### Male age and identity effects on egg-laying traits and Fledging Success

Male characteristics, including age, are increasingly shown to influence egg-laying traits in long-lived birds (Velando et al. 2006, Brommer and Rattiste 2008, Teplitsky et al. 2010, Auld et al. 2013), and accounting for male age can improve inference regarding female age effects (the focus of this study). Male and female Nazca boobies share incubation, brooding, and chick-provisioning (Anderson and Ricklefs 1992). Male Nazca boobies do not feed their mates, but may influence clutch phenotypes indirectly, by stimulating a female to reduce/increase investment into a given breeding attempt, for example (Dentressangle et al. 2008). We evaluated the role of male age and identity in shaping Nazca booby egg-laying traits and Fledging Success using the subset of breeding attempts involving a known-age, banded, male (sample size varied by response variable but was ≥ 7,662 records from ≥ 1,949 females and ≥ 2,226 males, Appendix S4). Non-breeding females are not associated with a partner’s age or identity, explaining the absence of Breeding Probability (evaluated only in females) at this analysis stage. The Punta Cevallos population is male-biased and females frequently change partners between breeding attempts, exchanging a male depleted by reproductive effort for a current non-breeder (Maness and Anderson 2007, 2008). This “mate rotation” helps to statistically decouple male versus female age and identity effects on breeding phenotypes. Male age was weakly correlated with female age (e.g., for Clutch Size: *r* = 0.29, d.f. = 7,927, *p* << 0.01; also see Fig. S1 in Tompkins and Anderson 2019) and was modelled as a three-level factor with young (≤ 9), middle-aged (10-15), and old (≥ 16) age classes (Tompkins and Anderson 2019).

GLMMs using the best-supported parameterization of female age effects from Age + Environment analyses were extended to include Male Identity as a random effect, categorical Male Age as a fixed factor, and an interaction between Female Age effects (continuous, threshold functions) and categorical Male Age. We also analyzed variation in Fledging Success at this stage, fitting a two-threshold age function with thresholds at ages 7 and 15 (following Tompkins and Anderson 2019). The Male Age by Female Age interactions allowed the slope estimates describing female age effects on breeding performance to vary with Male Age class (young, middle-aged, old). For Fledging Success, relative Breeding Date and A-egg Volume (both standardized within year) and Clutch Size (one vs. two eggs) were additional covariates, included to separate environmental and age effects on offspring production from those shaping correlated, and earlier-expressed, traits.

To evaluate trait variance associated with Male and Female Identity, three models with increasingly complex variance components (and the full fixed effect structure) were ranked using *el^p^^d PSIS-LOO* or *el^p^^d 10-fold CV*. The simplest model included only Breeding Season as a group-specific effect; next, Female Identity was added, and then Male Identity, all three as random intercept terms. Afterward, the best-supported random effect structure was retained and support for the interaction between Male and Female Age was evaluated from the span of the 95% BCI on interaction coefficients relative to zero.

#### Interactive effects of female age and environment on Breeding Probability, egg-laying traits and Fledging Success

Finally, we evaluated interactive effects of Female Age and environment on clutch-initiation traits and Fledging Success, using two analyses covering ages 3-12 (young to middle ages) and 11-20 (middle to old ages), respectively. Ages 3-12 include all of early-life improvement (Tompkins and Anderson 2019) and the earliest cohort – 1984 – reached age 12 in 1996, so that most breeding seasons included in the data span the full 3-12 age range. The final eight breeding seasons of the study (2010-2017) were used to evaluate the environmental dependence of late-life declines using data from known-age females aged 11-20. Breeding season and age ranges were chosen so that most years include the full 11-20 age range (exceptions: 2010 covers ages 11-18, 2011 and 2017 cover ages 11-19).

We evaluated interactions between age and environment using GLMMs retaining the best-supported parameterization of female age effects from Age + Environment analyses (for all traits, a threshold model) and including Female Identity and Breeding Season as random effects. Female Age effects appeared in pairwise interactions with our continuous measures of environmental quality. Interactions between aging rates and SSTA/SSTA^2^ or Nest Count were evaluated separately to avoid overfitting the GLMMs. The sign of interaction coefficients described how age-related improvement in early life and age-related declines in late life strengthen/weaken under environmental change, addressing the prediction that challenging environments should enhance performance discrepancies between young and middle-aged breeders and between middle-aged and old breeders. Parameter estimates for interaction coefficients were evaluated by the span of the 95% BCI relative to zero.

Any influence of age by environment interactions on population dynamics will act through offspring production. We ran two versions of the Fledging Success “Age by Environment” interaction models, one with, and one without, controlling the effects of traits that are expressed earlier in the breeding season. When these early traits are excluded from the models, coefficient estimates convey the total effects of age by environment interactions on Fledging Success, including those carried through Clutch Size, Breeding Date, or A-egg Volume and their relationship with reproductive success.

## RESULTS

### Influence of female age on egg-laying traits

Middle-aged female Nazca boobies laid clutches earlier in the breeding season, had larger A-egg Volumes, and had higher probabilities of laying a two-egg clutch than young or very old females (Fig. 1). The age-dependence of egg-laying traits was supported statistically: candidate models excluding age effects had much lower out-of-sample predictive accuracy than models including age (Table 2, full results in Appendix S8: Table S1). Two-threshold age functions outperformed linear, quadratic, and single-threshold options for all egg-laying traits (Table 2).

**Figure 1.**
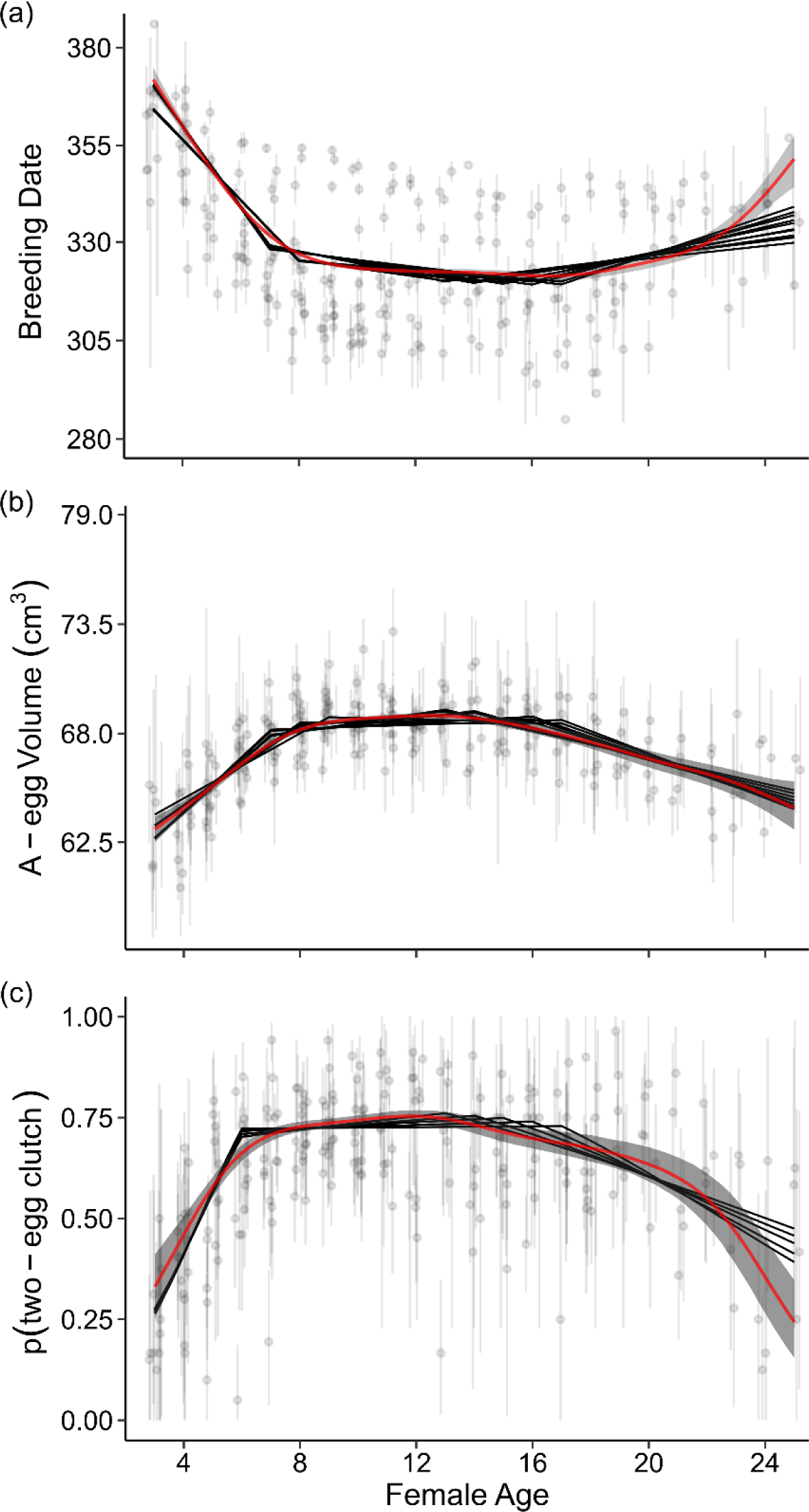
Female age-specific Breeding Date. (a), A-egg Volume (b), and probability of laying a two-egg clutch (c), in Nazca boobies. Breeding Dates in extended Julian dates that span a two-year period. Red lines are model predictions from Age + Environment GAMMs; shaded regions cover 95% BCIs. Black lines are model predictions from top threshold models for each trait.

**Table 2.** Expected log predictive densities (*el^p^^d*) from 10-fold cross-validation support female age effects on Breeding Date, Clutch Size, and A-egg Volume in Nazca boobies. Δelp^*d* gives distance from the best-supported GLMM (in bold), with negative Δ*el^p^^d* indicating relatively poor out-of-sample predictive ability. Model rankings compare alternative parameterizations of age effects, including a non-parametric smooth function (“s(F Age)”) and single-threshold (“One T”; only the best-supported candidate is shown) or two-threshold functions (“Two T”; all candidates whose performance overlaps the best-supported GLMM are shown). Model-estimated aging rates for early, prime age, and late periods are given for two-threshold models.

| Age model | $\Delta \ell \hat{p} d_{10\text{-fold } CV} (SE)$ | Slope 1 [95% BCI] | Slope 2 [95% BCI] | Slope 3 [95% BCI] |
| --- | --- | --- | --- | --- |
| **Breeding Date** |  |  |  |  |
| s(F Age) | 61.6 (21.5) |  |  |  |
| **Two T (7, 15)** | **0.0** | **-4.24 [-4.43, -4.06]** | **-0.48 [-0.54, -0.41]** | **0.64 [0.52, 0.75]** |
| Two T (8, 17) | -2.7 (23.7) | -3.19 [-3.32, -3.06] | -0.24 [-0.29, -0.18] | 0.91 [0.75, 1.07] |
| Two T (7, 13) | -7.5 (18.6) | -4.14 [-4.34, -3.95] | -0.63 [-0.71, -0.55] | 0.38 [0.30, 0.47] |
| Two T (8, 14) | -12.5 (23.6) | -3.16 [-3.29, -3.02] | -0.37 [-0.45, -0.29] | 0.44 [0.34, 0.54] |
| Two T (7, 17) | -15.1 (18.6) | -4.31 [-4.50, -4.13] | -0.37 [-0.42, -0.31] | 1.02 [0.86, 1.17] |
| Two T (7, 16) | -16.3 (18.7) | -4.28 [-4.47, -4.08] | -0.42 [-0.48, -0.36] | 0.81 [0.67, 0.94] |
| Two T (7, 14) | -25.2 (18.4) | -4.19 [-4.39, -4.00] | -0.55 [-0.62, -0.48] | 0.50 [0.40, 0.60] |
| Two T (8, 16) | -25.3 (23.6) | -3.18 [-3.31, -3.04] | -0.28 [-0.34, -0.21] | 0.71 [0.58, 0.84] |
| Two T (8, 15) | -35.6 (23.5) | -3.17 [-3.30, -3.04] | -0.32 [-0.39, -0.25] | 0.56 [0.44, 0.67] |
| Two T (8, 13) | -48.1 (24.0) | -3.13 [-3.27, -2.99] | -0.43 [-0.52, -0.34] | 0.33 [0.24, 0.42] |
| One T (8) | -101.4 (26.5) |  |  |  |
| Age + Age<sup>2</sup> | -208.4 (30.4) |  |  |  |
| Age | -1,018.9 (49.9) |  |  |  |
| No Age | -1,288.8 (57.5) |  |  |  |
| **A-egg Volume** |  |  |  |  |
| s(F Age) | -48.1 (39.7) |  |  |  |
| **Two T (7, 13)** | **0.0** | **2.27 [2.07, 2.48]** | **0.39 [0.31, 0.48]** | **-0.62 [-0.71, -0.53]** |
| Two T (8, 13) | -46.9 (39.8) | 1.77 [1.62, 1.91] | 0.27 [0.17, 0.36] | -0.59 [-0.68, -0.50] |
| Two T (9, 16) | -56.9 (39.1) | 1.45 [1.34, 1.56] | -0.07 [-0.15, 0.01] | -0.78 [-0.91, -0.63] |
| Two T (8, 14) | -57.4 (41.5) | 1.80 [1.65, 1.94] | 0.19 [0.10, 0.27] | -0.69 [-0.79, -0.58] |
| Two T (8, 17) | -62.5 (37.2) | 1.84 [1.70, 1.99] | -0.01 [-0.07, 0.06] | -0.93 [-1.09, -0.77] |
| Two T (8, 16) | -72.6 (50.1) | 1.83 [1.69, 1.98] | 0.05 [-0.03, 0.12] | -0.84 [-0.99, -0.71] |
| Two T (7, 16) | -75.8 (52.6) | 2.41 [2.21, 2.62] | 0.14 [0.08, 0.21] | -0.90 [-1.05, -0.76] |
| Two T (7, 17) | -77.6 (43.2) | 2.43 [2.22, 2.64] | 0.09 [0.02, 0.15] | -1.00 [-1.16, -0.83] |
| Two T (7, 14) | -82.9 (43.9) | 2.33 [2.13, 2.52] | 0.30 [0.22, 0.38] | -0.73 [-0.83, -0.63] |
| One T (7) | -100.8 (18.6) |  |  |  |
| Age + Age<sup>2</sup> | -164.5 (40.9) |  |  |  |
| Age | -426.4 (51.5) |  |  |  |
| No Age | -456.4 (46.7) |  |  |  |
| **Clutch Size** |  |  |  |  |
| s(F Age) | 4.4 (6.7) |  |  |  |
| **Two T (6, 13)** | **0.0** | **6.06 [4.99, 7.07]** | **0.43 [0.20, 0.65]** | **-1.05 [-1.30, -0.80]** |
| Two T (6, 17) | -5.1 (6.9) | 6.62 [5.63, 7.64] | 0.03 [-0.12, 0.18] | -1.79 [-2.30, -1.23] |
| Two T (6, 14) | -5.9 (5.6) | 6.22 [5.20, 7.24] | 0.29 [0.09, 0.48] | -1.18 [-1.50, -0.87] |
| Two T (6, 15) | -9.3 (6.0) | 6.37 [5.39, 7.40] | 0.19 [0.01, 0.36] | -1.34 [-1.69, -0.99] |
| Two T (6, 16) | -11.9 (6.5) | 6.48 [5.46, 7.53] | 0.11 [-0.06, 0.27] | -1.55 [-1.97, -1.12] |
| Two T (7, 17) | -13.3 (7.9) | 3.98 [3.36, 4.62] | -0.10 [-0.26, 0.06] | -1.64 [-2.18, -1.13] |
| One T (6) | -28.4 (8.7) |  |  |  |
| Age + Age<sup>2</sup> | -30.5 (9.9) |  |  |  |
| Age | -117.9 (16.3) |  |  |  |
| No Age | -120.5 (16.4) |  |  |  |

We considered candidate models to have similar explanatory ability whenever the *el^p^^d* difference between them was small relative to its SE. For all egg-laying traits, there was substantial model selection uncertainty regarding the threshold age dividing prime age from late life (models with the second threshold age at 13-17 years of age have Δ*el^p^^d* values less than 2 times the SE; Table 2). However, despite uncertainty in the onset of performance declines, all top models estimate late-life aging rates that are distinct from zero, consistent with senescence (Table 2). For Breeding Date (and, to a lesser extent, Clutch Size), the best-performing two-threshold model failed to represent accelerating late-life performance decline (Fig. 1) and the GAMMs modelling each response variable as a smooth function of Female Age had greater out-of-sample predictive accuracy (Table 2).

From Selective Disappearance GLMMs, our proxy for lifespan did not covary with Breeding Date (β ALA = −0.03 [95% BCI: −0.07, 0.02]), A-egg Volume (β ALA = 0.00 [95% BCI: −0.06, 0.07] or Clutch Size (β ALA = 0.02 [95% BCI: −0.08, 0.12]), suggesting little opportunity for compositional change in an aging cohort to obscure or artificially enhance estimated age functions.

### Influence of male age and identity on egg-laying traits and Fledging Success

Repeatable elements of each pair member’s phenotype contributed to Breeding Date (supported by *el^p^^d PSIS-LOO* model rankings; Table 3). Repeatabilities for each sex were calculated as intra-class correlation coefficients (ICC), the proportion of variance not accounted for by fixed effects in the model (values 0 to 1). Male repeatability for Breeding Date was very low (ICC = 0.06, Table 3), and smaller in magnitude than female repeatability (ICC = 0.19, Table 3).

**Table 3.**
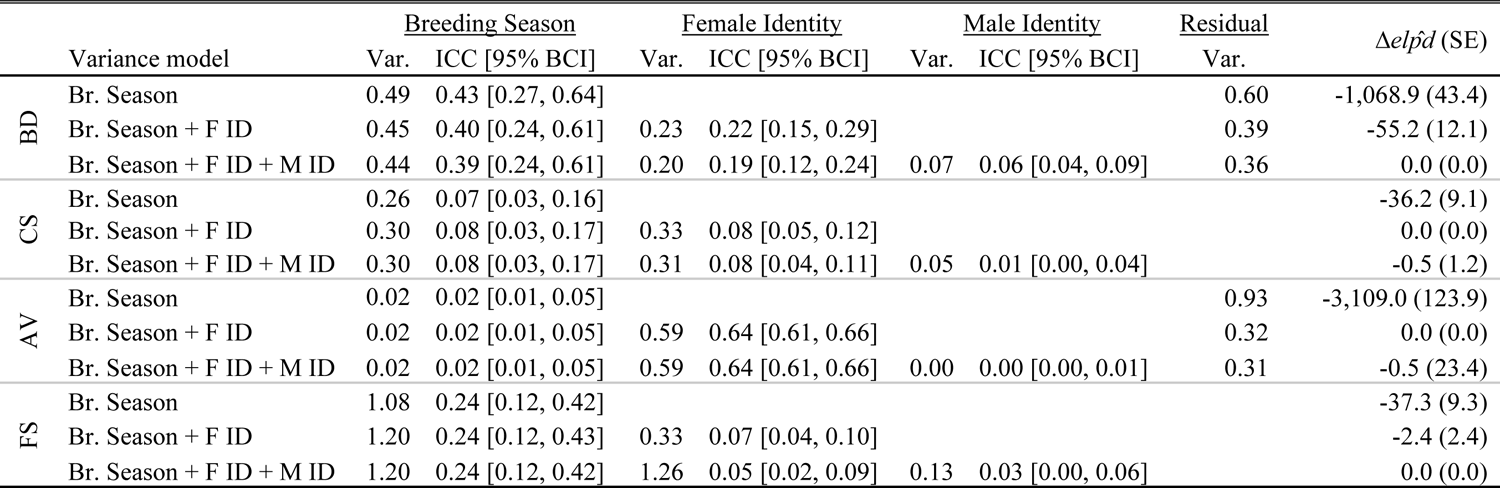
Model selection results evaluating the influence of Male (M ID) and Female Identity (F ID) on Breeding Date (BD), Clutch Size (CS), A-egg Volume (AV), and Fledging Success (FS) in Nazca boobies. Models were ranked by 10-fold cross validation (A-egg Volume) or PSIS-LOO (all others), and negative Δ*el^p^^d* (relative to the best-supported model) indicate worse model predictive performance. Models included environmental and other fixed effects, held constant across candidate variance component configurations.

| | Variance model | <u>Breeding Season</u> | | <u>Female Identity</u> | | <u>Male Identity</u> | | <u>Residual</u> | $\Delta\hat{elpd}$ (SE) |
| --- | --- | --- | --- | --- | --- | --- | --- | --- | --- |
|  |  | Var. | ICC [95% BCI] | Var. | ICC [95% BCI] | Var. | ICC [95% BCI] | Var. |  |
| BD | Br. Season | 0.49 | 0.43 [0.27, 0.64] |  |  |  |  | 0.60 | -1,068.9 (43.4) |
|  | Br. Season + F ID | 0.45 | 0.40 [0.24, 0.61] | 0.23 | 0.22 [0.15, 0.29] |  |  | 0.39 | -55.2 (12.1) |
|  | Br. Season + F ID + M ID | 0.44 | 0.39 [0.24, 0.61] | 0.20 | 0.19 [0.12, 0.24] | 0.07 | 0.06 [0.04, 0.09] | 0.36 | 0.0 (0.0) |
| CS | Br. Season | 0.26 | 0.07 [0.03, 0.16] |  |  |  |  |  | -36.2 (9.1) |
|  | Br. Season + F ID | 0.30 | 0.08 [0.03, 0.17] | 0.33 | 0.08 [0.05, 0.12] |  |  |  | 0.0 (0.0) |
|  | Br. Season + F ID + M ID | 0.30 | 0.08 [0.03, 0.17] | 0.31 | 0.08 [0.04, 0.11] | 0.05 | 0.01 [0.00, 0.04] |  | -0.5 (1.2) |
| AV | Br. Season | 0.02 | 0.02 [0.01, 0.05] |  |  |  |  | 0.93 | -3,109.0 (123.9) |
|  | Br. Season + F ID | 0.02 | 0.02 [0.01, 0.05] | 0.59 | 0.64 [0.61, 0.66] |  |  | 0.32 | 0.0 (0.0) |
|  | Br. Season + F ID + M ID | 0.02 | 0.02 [0.01, 0.05] | 0.59 | 0.64 [0.61, 0.66] | 0.00 | 0.00 [0.00, 0.01] | 0.31 | -0.5 (23.4) |
| FS | Br. Season | 1.08 | 0.24 [0.12, 0.42] |  |  |  |  |  | -37.3 (9.3) |
|  | Br. Season + F ID | 1.20 | 0.24 [0.12, 0.43] | 0.33 | 0.07 [0.04, 0.10] |  |  |  | -2.4 (2.4) |
|  | Br. Season + F ID + M ID | 1.20 | 0.24 [0.12, 0.42] | 1.26 | 0.05 [0.02, 0.09] | 0.13 | 0.03 [0.00, 0.06] |  | 0.0 (0.0) |

Female Identity explained variation in Clutch Size (ICC = 0.08) and A-egg Volume (ICC = 0.64; Table 3). Male repeatabilities were small for both traits (≤ 0.01), and the inclusion of Male Identity did not improve model performance (Table 3), emphasizing that differences among females, not males, are most relevant to expression of Clutch Size and A-egg Volume.

The model partitioning variation in Fledging Success by Male Identity, Female Identity, and Breeding Season had the best predictive accuracy, but was not significantly better than a simpler model excluding Male Identity effects (Table 3) and mixed poorly with respect to the identity terms, requiring double the iterations to reach the target effective sample size as simpler models. Estimated repeatabilities for both sexes under this model were low (male ICC = 0.03, female ICC = 0.05, Table 3); while we fail to disentangle male and female identity effects for this trait, these effects are small.

Male and female ages interacted to influence Breeding Date. Old females paired with young males experienced delays in Breeding Date with advancing age (aging rate = 0.85 [95% BCI: 0.54, 1.18]); females paired with middle-aged (slope = 0.23 [95% BCI: −0.01, 0.45] or old partners (slope = 0.12 [95% BCI: −0.25, 0.50]) did not (Fig. 2, coefficient estimates in Appendix S8: Table S3). Male age influenced Clutch Size and Fledging Success but did not interact with Female Age (the 95% BCI of all interaction coefficients included zero). Pairing with a young partner rather than a middle-aged one reduced the probability of laying a two-egg clutch by 0.02 (β = −0.21 [95% BCI −0.34, −0.08]), holding age at 8 years and all other predictors in the model at their mean value. No additional disadvantage of pairing with a young male, beyond that carried by the positive influence of Clutch Size on reproductive success, was found for Fledging Success (β = −0.10 [95% BCI: −0.23, 0.02]). Breeding with an old male rather than a middle-aged partner reduced Fledging Success (range 0-1) by 0.08 (β = −0.40 [95% BCI: −0.59, −0.20]) but did not affect Clutch Size or A-egg Volume (coefficient estimates in Appendix S8: Table S3).

**Figure 2.**
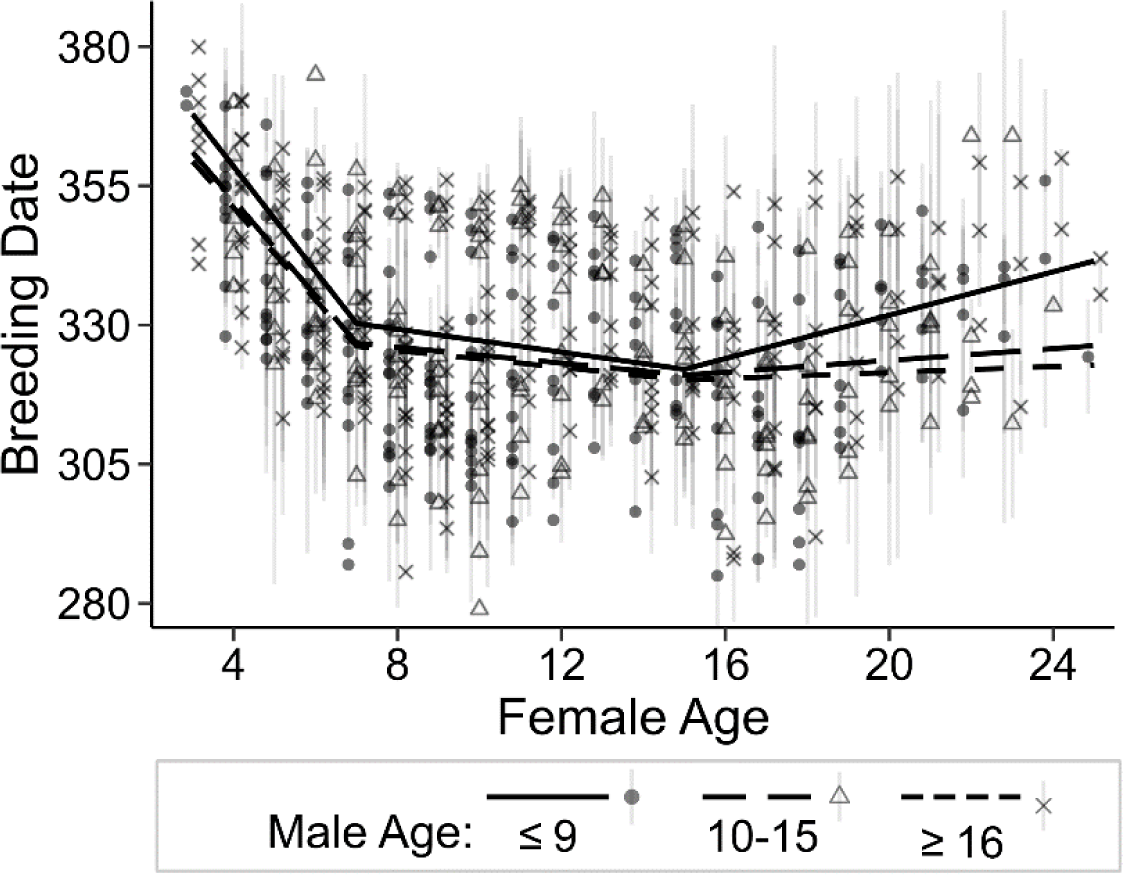
Male Age interacts with Female Age to determine Breeding Date in Nazca boobies. Breeding Dates are in extended Julian dates. Model predictions (lines) and 95% BCI are from a GLMM modeling Female Age as a two-threshold function interacting with categorical Male Age. Points and intervals are means and 95% CI of the raw data binned by Female Age and categorical Male Age.

### Influence of environmental variation on egg-laying traits

Clutch Size covaried positively with SSTA, our proxy for yearly differences in the ENSO (β = 0.30 [95% BCI: 0.10, 0.49]; model coefficient estimates in Appendix S8: Table S2). Large clutch sizes under positive SSTA signaled that El Niño-like conditions are a relatively resource-rich environment during egg-laying, and La Niña-like conditions are a relatively poor one. Consistent with this interpretation, in Nov.-Jan. the probability that a sampled Nazca booby regurgitated a food item increased marginally with SSTA (β = 0.30 [95% CI: −0.04, 0.63]). Clutch Size also covaried positively with Nest Count, increasing with the number of breeding pairs (β = 0.27 [95% BCI: 0.12, 0.43]). In contrast, Breeding Date and A-egg Volume were not affected by yearly variation in either continuous measure of environmental quality (Appendix S8: Table S2). Fish Phase did not alter egg-laying traits (Appendix S8: Table S2).

Posterior predictive checks of Age + Environment models for Breeding Date suggested the model fit the data poorly in two regards: random intercepts for Breeding Seasons 2013-2016 were much larger than expected under the assumption that all Breeding Season effects are drawn from the same Gaussian distribution, and the variance in Breeding Date was not constant across seasons. However, after adjustments to control the lack of fit, our results with respect to Female Age, SSTA, and Nest Count were unchanged (details in Appendices S9-S10).

### Interactive effects of female age and environment on Breeding Probability, egg-laying traits and Fledging Success

#### Age span 3-12

In the first half of the lifespan, Female Age interacted with interannual differences in the ENSO to explain variation in Breeding Probability, Breeding Date, and Clutch Size, but not A-egg Volume or Fledging Success. As expected, the Breeding Probability of young Nazca boobies responded most strongly to variation in SSTA (Fig. 3a): young females drastically reduced their breeding probability in cool La Niña-like conditions relative to years with high SSTA (e.g., a reduction in Breeding Probability from 0.40 to 0.15 at age 4), while prime age birds did not (the corresponding reduction in Breeding Probability is only 0.99 to 0.94 at age 10). The same pattern was observed for Breeding Date (Fig. 3b): young birds laid earlier when surface waters were relatively warm while middle-aged birds showed no change in Breeding Date (coefficient estimates in Appendix S8: Table S4). However, the forms of age by environment interactions were not consistent across traits. Poor la Niña-like conditions showed the shallowest aging trajectories for Clutch Size (Fig. 3c), and the same trend was predicted for A-egg Volume, although the 95% BCI of the relevant coefficient spanned zero (Appendix S8: Table S4).

**Figure 3.**
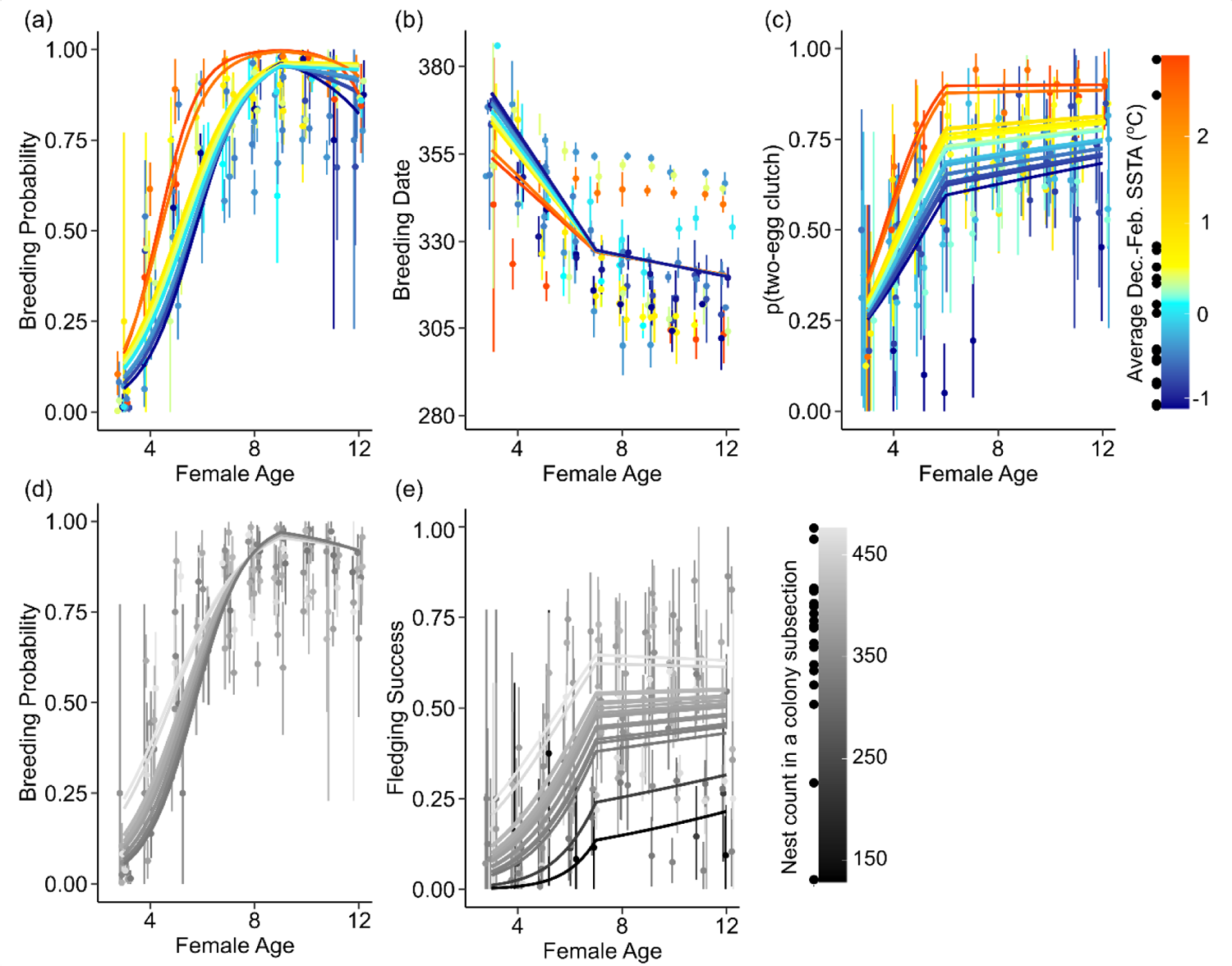
Age interacts with continuous measures of environmental quality to influence Breeding Probability (a,d), Breeding Date (b), Clutch Size (c), and Fledging Success (e, a marginal effect) in young and middle-aged female Nazca boobies (ages 3-12). The model-predicted age by environment interaction is shown at each value of SSTA (a,b,c) or Nest Count (d,c) present in the data (lines; also see points along legend gradients). Points and intervals are means and 95% CI of the raw data binned by Female Age and Breeding Season. Breeding Dates are in extended Julian dates.

In the first half of the lifespan, Female Age interacted with Nest Count to explain variation in Breeding Probability and Fledging Success but not Breeding Date, A-egg Volume, or Clutch Size (Appendix S8: Table S5). Breeding Probability followed expectations: young females, more than middle-aged ones, reduced their probability of initiating a clutch in years with low Nest Count, increasing age-related differences in Breeding Probability in a poor environment (Appendix S8: Table S5). The coefficient estimate describing the effect of Nest Count on early life improvement in Fledging Success implied shrinking age-related changes with increasing Nest Count, a marginal effect (β = −0.80 [-1.60, 0.02], Appendix S8: Table S5), and only appeared when traits that are expressed earlier were omitted from the model. However, on the probability scale (Fig. 3d), age differences in Fledging Success are relatively low, not high, at the lowest Nest Counts because age trajectories are compressed close to the lower boundary of zero.

#### Age span 11-20

Late in life (out to age 20), interannual variation in SSTA interacted with Female Age to influence Breeding Date, but not the other reproductive traits (Appendix S8: Table S6). In the last eight years of the study, El Niño-like conditions delayed breeding, particularly in the oldest females (Fig. 4a, Appendix S8: Table S6). Late in life, Nest Count interacted with Female Age to influence Fledging Success, but not Breeding Probability or egg-laying traits (Appendix S8: Table S7). Age-related changes in Fledging Success before the late-life threshold age (age 15) were influenced by Nest Count (β = 0.59 [95% BCI: 0.06, 1.12]): when clutch initiations were high, performance was constant through middle age, but when clutch initiations were low, performance declined from age 11 (the youngest age included) onward (Fig. 4b). After the threshold age, aging rate was not affected by Nest Count (β = −0.92 [95% BCI: −2.04, 0.23]; thus Nest Count influences late-life aging by altering the onset of late-life declines, and not by affecting the steepness of senescence for the oldest age classes.

**Figure 4.**
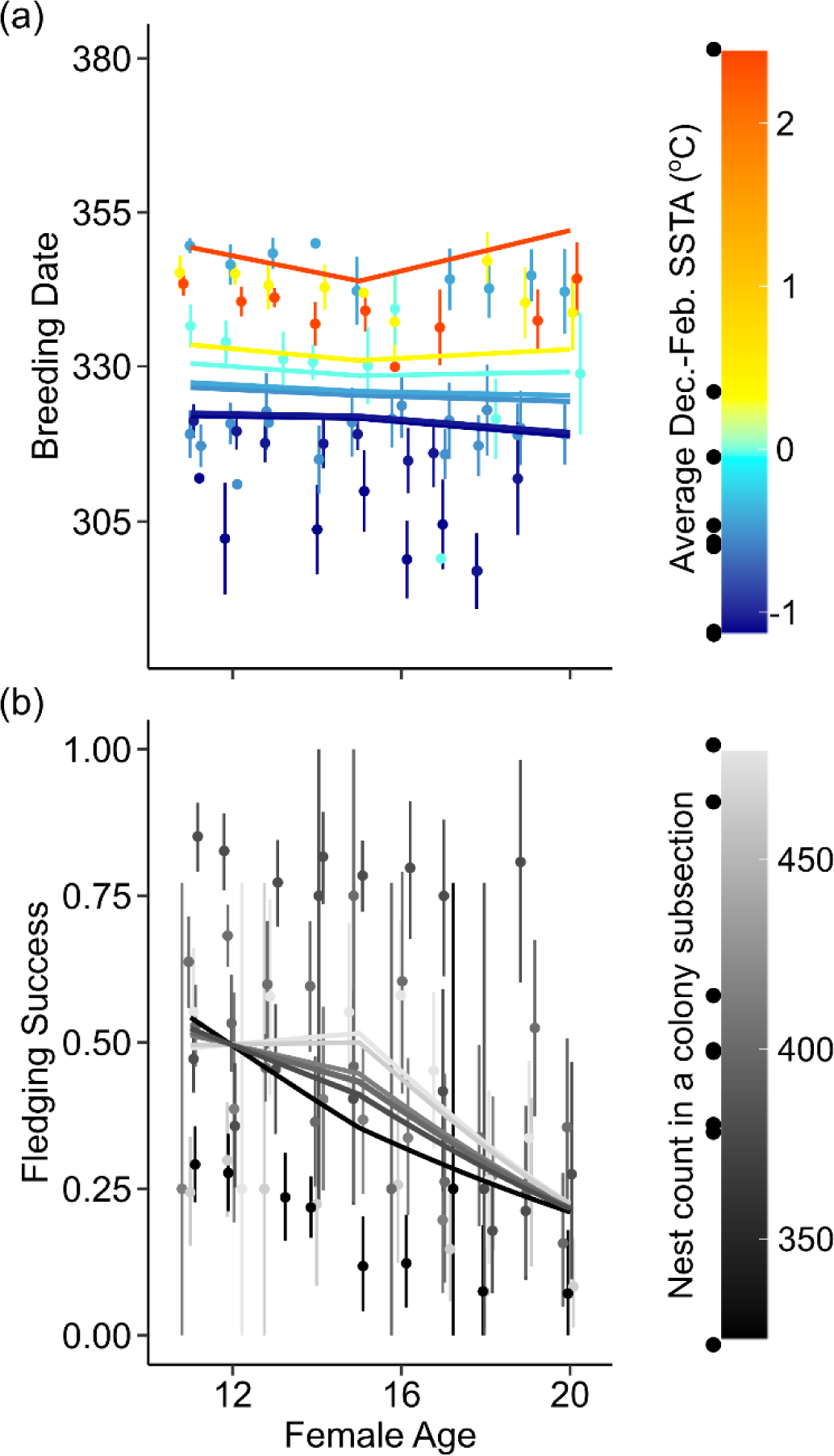
Late in life, aging patterns depend on SSTA for Breeding Date (a) and on Nest Count (b) for Fledging Success. Model-predicted age by environment interactions are shown at each value of SSTA/Nest Count present in the data (lines; also see points along legend gradients). Points and intervals are means and 95% CI of the raw data binned by Female Age and Breeding Season. Breeding Dates in extended Julian dates.

## DISCUSSION

Middle-aged females initiated clutches earlier, had higher probabilities of laying a second egg, and laid larger eggs than young or old Nazca boobies. These patterns echo early-life improvements and late-life declines in Breeding Probability and Fledging Success (Tompkins et al. 2017, Tompkins and Anderson 2019), emphasizing that age is a crucial factor structuring variation in fitness-related traits in this population. Age effects were driven by processes acting within individuals; selective disappearance of low-quality phenotypes cannot explain them.

Young and old boobies were predicted to perform especially poorly in challenging environments (Ratcliffe et al. 1998, Laaksonen et al. 2002). In early life, responses to environmental quality (ENSO variation, Nest Count) depended on age for many traits/measures of environment: age-related differences in performance shrank with increasing environmental quality for Breeding Probability and Breeding Date, but not for Clutch Size or Fledging Success. In late life, delays in Breeding Date and declines in Fledging Success depended on ENSO status and Nest Count, respectively, confirming environmental effects on age-specific patterns across the entire lifespan. We first discuss the evidence for senescence in reproductive performance across the breeding cycle, and then confront the diversity of age by environment interactions revealed in this study with respect to inferred “good” and “poor” environments for breeding.

### Early-life improvement

Young female Nazca boobies performed worse for all traits examined than did middle-aged females. Our data do not allow us to distinguish the effects of reproductive restraint from those of inexperience with breeding and/or foraging. However, the non-linear form of aging patterns (performance increased steeply in young breeders before leveling off) suggests that acquisition of skills during the first few breeding attempts plays some role in forming observed patterns (Curio 1983). Familiarity and synchronization with the male partner may also be important, especially for Breeding Date, which advanced (progressively more slowly) across much of the adult lifespan (Table 2). Although female Nazca boobies rotate mates frequently between breeding seasons (Maness and Anderson 2007, 2008), in doing so they may return to a previous partner or engage with a familiar neighbor (breeding site fidelity is high; Huyvaert and Anderson 2004), contributing to continued improvement through middle age. Selective disappearance of low-quality phenotypes cannot explain these early-life improvements. We find no support for the progressive removal of low-quality individuals from an aging cohort with respect to Breeding Date, Clutch Size, or A-egg Volume (this study, Townsend and Anderson 2007a). Differences in phenotypic quality do exist among female Nazca boobies (Townsend and Anderson 2007a,b, Tompkins and Anderson 2019), but are uncorrelated with lifespan.

### Late-life decline

Actuarial and reproductive senescence are common in the wild, but our understanding of the causes and consequences of variation in senescence within populations – by environments, and across different components of reproduction and self-maintenance – remains incomplete (Nussey et al. 2013, Lemaître and Gaillard 2017). Beginning around age 15, Nazca booby females start modest declines in breeding probability and relatively steep declines in annual survival probability and fledging success (Tompkins and Anderson 2019). Late in life, Nazca booby females also initiate their clutch relatively late in the season and lay smaller clutches and A-eggs (this study), adding clutch initiation phenotypes to the list of traits showing senescence in Nazca boobies.

Early in the breeding cycle, late-life declines in egg-laying traits are relatively minor: A-egg Volumes shrink ∼4.5 cm^3^ over 10 years (less than 1 s.d., Fig. 1, Table 2) while steep age-related breeding delays and reductions in Clutch Size are experienced only at the oldest ages (≥ age 20, Fig. 1). These egg-laying traits affect offspring production: Fledging Success improves with early breeding (β = −0.13 [95% BCI: −0.19, −0.07]), laying a two-egg clutch (β = 0.67 [95% BCI: 0.55, 0.80]), and laying A-eggs of larger volume (β = 0.08 [95% BCI: 0.02, 0.14], Appendix S8: Table S3). Thus, while age-related changes in egg-laying phenotypes are weak, performance integrated across all reproductive stages (including those not examined here) results in strong senescent decline in Fledging Success: in average conditions, Fledging Success is halved between the ages of 12 and 20 (Tompkins and Anderson 2019). Looking across traits, late-life declines, relative to peak performance, also are stronger for offspring production versus egg-volume and laying date in blue-footed boobies (*Sula nebouxii*; Beamonte-Barrientos et al. 2010, Kim et al. 2011) and wandering albatrosses (*Diomedea exulans*; Froy et al. 2013; but see Ratcliffe et al. 1998 for the opposite pattern in great skuas). As proposed for mammals (Nussey et al. 2009), physiological deterioration associated with senescence may constrain the expression of reproductive traits in step with their energetic cost, explaining stronger senescence in traits (like Fledging Success) dependent on parental care across the breeding cycle.

### Interactions between age and environment

Young individuals, constrained by inexperience or practicing reproductive restraint, were expected to perform particularly poorly in a resource-poor environment. This expectation was met for Breeding Probability and Breeding Date in Nazca boobies. Results of the current study (based on Clutch Size and an index of food availability) indicated that El Niño-like conditions (positive SSTA) represented a relatively resource-rich environment during egg-laying and La Niña-like conditions (negative SSTA) a poor one. Relative to prime-age birds, young females were particularly likely to skip breeding or delay breeding for the first time (our analysis does not distinguish these events) in a poor environment (indicated by a negative SSTA or low Nest Count; Fig. 3a,d). Young birds also initiated their clutches particularly late in the breeding season in poor La-Niña like conditions (Fig. 3b). In contrast with Breeding Date and Breeding Probability, environment by age interactions on Clutch Size and Fledging Success did not follow the expected pattern. The Clutch Size of middle-aged birds increased faster with SSTA than that of young birds, enhancing age-related differences when sea surface temperatures were relatively warm (Fig. 3c). For Fledging Success, a marginal, negative, coefficient estimate (β = −0.80 [95% BCI: −1.60, 0.02]) describing the effect of Nest Count on early-life improvement implied shrinking age effects as the environment improved. However, variation in Fledging Success was modelled on the logit scale. When the age by Nest Count interaction is translated back to the biologically-relevant probability scale the opposite pattern appears (Fig. 3e): in a very poor environment, Fledging Success is low at all ages (close to the boundary of zero), compressing the differences between young and middle-aged females.

Across studies, the emerging picture suggests that early-life improvements in breeding performance often vary with environmental conditions (but see Vieyra et al. 2009 and Pardo et al. 2013 for exceptions). The expected pattern – accentuated age effects on breeding performance in a resource-poor environment – has been observed for fecundity and pregnancy rate in red deer (Clutton-Brock et al. 1987, Bonenfant et al. 2002), clutch size in Tengmalm’s owls (*Aegolius funereus*; Laaksonen et al. 2002), offspring production in seabirds (Sydeman et al. 1991, Bunce et al. 2005), and is implied by higher and less variable fecundity in prime age versus young ungulates (Bjørkvoll et al. 2016, reviewed in Gaillard et al. 2000). Breeding Probability and Breeding Date followed the expected pattern in female Nazca boobies.

However, our study shows that the form taken by age-environment interactions can vary considerably, even within a population. Diminished – not enhanced – age effects on breeding performance in a resource-poor environment have been observed for clutch size, egg volume, and hatching success in Audouin’s gulls (Oro et al. 2014), for breeding date and egg volume in Scopoli’s shearwaters (Hernández et al. 2015), and now for Clutch Size and Fledging Success in Nazca boobies.

Only breeding individuals express Clutch Size and Fledging Success phenotypes in a given year. Thus, harsh environments may shrink age-related differences in Clutch Size, for example, if young individuals in relatively poor condition (due to inexperience or low quality) fail to breed (e.g., Weimerskirch 1992, Becker and Bradley 2007). This explanation has been proposed by Oro et al. (2014) and Hernández et al. (2015) to explain larger age-related differences in breeding performance under high resource availability, although neither study explicitly evaluated breeding participation. Middle-aged Nazca boobies rarely skip breeding, while young birds are increasingly likely to skip in a poor versus a good environment (Fig. 3a,d). Thus, the composition of young breeders in a good environment may include a relatively high proportion of poor quality or inexperienced Nazca boobies (females breed for the first time at 3-7 years of age; Champagnon et al. 2018), contributing to the unexpected patterns for Clutch Size and Fledging Success. The relationship of physical condition (or previous experience) to environment-dependent breeding probabilities in young females has yet to be examined.

The life history syndrome of Nazca boobies (high annual adult survival, delayed recruitment, one offspring raised per breeding season; Anderson 1990, Townsend and Anderson 2007b, Apanius et al. 2008, Maness and Anderson 2013) suggests that they should prioritize resource allocation toward self-maintenance over reproduction (compared to short-lived organisms; Stearns 1992), flexibly adjusting reproductive effort and cost in response to environmental conditions (Erikstad et al. 1998, Townsend and Anderson 2007). Stronger reproductive responses to environmental variation by middle-aged individuals and the expansion of age effects in benign environments, as we observe for Clutch Size and Fledging Success, may be the outcome of selection for low reproductive investment (even by prime-age individuals) in a harsh environment to preserve future opportunities. Looking across traits, our results suggest that young Nazca boobies, facing interannual environmental variation, moderate reproductive effort at an earlier stage (breeding probability) than middle-aged birds (clutch size and fledging success), contributing to the trait-dependence of age-environment interactions. This pattern is consistent with a lower average condition in young adults (e.g., Weimerskirch 1992, Barbraud and Weimerskirch 2005, Becker and Bradley 2007) and sequential, condition-dependent, decisions to breed or not breed (Drent and Daan 1980, Weimerskirch 1992), lay a large or small clutch (e.g., in Nazca boobies; Clifford and Anderson 2001,2002) or invest in providing food for offspring or one’s self.

The senescence and terminal restraint hypotheses (McNamara et al. 2009) each predict that, in poor environments, old individuals will show greater performance reductions than middle-aged individuals because of somatic degradation and/or low investment in reproduction. The last eight years of our study were used to evaluate environmental effects on aging patterns across middle to old ages in Nazca boobies. For Fledging Success, and following predictions, age-related performance declines began at a younger age when Nest Counts were low than when they were high (Fig. 4b). Looking across the full lifespan, changes in Fledging Success with age were compressed during poor environmental conditions (low Nest Count) across young to middle ages but enhanced across middle to old ages. This suggests that age-dependent responses to environmental variation may take on different forms for early-life improvement vs. senescent decline, but we note that the last eight years of the study exclude the lowest Nest Counts appearing in the full data (Fig. 3c vs. Fig. 4b). Age effects on Fledging Success in the second half of the lifespan may be limited in a very poor environment (as in the first half of the lifespan), but the range of conditions for which we have data from known-age old birds is still too narrow to test this idea.

Age-dependent responses to environmental variability also extended into old age for Breeding Date with respect to variation in SSTA. Contrary to expectations, middle-aged and old individuals maintained similar Breeding Dates in “poor” La Niña-like conditions, while “good” El Niño-like conditions were associated with the strongest age effects on performance.

However, attributing variation in the pattern of late-life age-dependence for Breeding Date solely to SSTA might be premature. Reproductive data from females aged 11-20 were available in eight years; in four of those years (2013-2016), breeding exhibited an unusual delay (by about a month, Appendix S9: Fig. S2), suggesting the existence of at least one additional factor exerting a strong influence on the timing of laying (only one of the delayed seasons, 2015, occurred during a strong El Niño warm event). Regardless of the environmental cause(s), variation in aging rate was apparent across the eight breeding seasons analyzed, such that performance declines with age were limited to certain environments for this trait.

To conclude this section, we used 20 years of longitudinal data on the reproductive phenotypes of Nazca boobies to evaluate interactive effects of age and environment on Breeding Probability, egg-laying traits and Fledging Success across the lifespan, revealing complex age-, environment-, and trait-dependent patterns of performance. Following predictions from life history theory, studies evaluating age by environment interactions affecting adult survival often find that the survival of young and/or old individuals is reduced to a greater extent than that of prime-age animals in poor environments (e.g., Coulson et al. 2001, Tavecchia et al. 2005, Barbraud and Weimerskirch 2005, Nevoux et al. 2007, Pardo et al. 2013). Our study, and others (e.g., Oro et al. 2014), confirm that breeding responses to environmental conditions can also be age-dependent across the entire lifespan, but – in contrast with survival – the resulting patterns of trait expression can be complex, differing across populations and, within populations, across traits. Adult survival is high, relatively stable, and closely related to fitness in species with moderate to long lifespans (Gaillard et al. 2000, Doherty et al. 2004); resource allocation trade-offs may be less likely to complicate optimal patterns of survival probabilities with respect to age and environment. Proxies for environmental quality used in this study did not measure prey availability directly, which may have introduced error and complicated interpretation relative to simple predictions based on aging theory, but also lead to improved insight into breeding responses to the ENSO, a phenomenon of global importance and increasing in frequency in our changing climate (Yeh et al. 2009, Cai et al. 2014).

### Partner’s influence on egg-laying traits and Fledging Success

While our focus was on reproductive aging in female boobies, we also found that a breeding partner’s age influenced a female’s egg-laying traits and Fledging Success. Breeding with a young male reduced the probability of laying a two-egg clutch, and delayed laying relative to pairing with a middle-aged mate. The same trend was observed for Fledging Success but was supported only weakly. Timing of clutch initiation is increasingly shown to depend on male as well as female phenotypes in long-lived birds (Laaksonen et al. 2002, Brommer and Rattiste 2008, Teplitsky et al. 2010, Auld and Charmantier 2011). Egg volume may also respond to male characteristics (e.g., in blue-footed boobies, Dentressangle et al. 2008), while expression of clutch size is usually sex-limited. Our results are remarkable for finding effects of Male Age on Breeding Date and Clutch Size (and also for Fledging Success, where breeding with old males depressed performance), given that male Nazca boobies do not feed their mates. The influence of Male Age on egg-laying traits is apparently indirect. Females may reduce investment into clutches when paired with a mate perceived to be in poor condition (low quality, inexperienced, or perhaps experiencing senescence), as in blue-footed boobies (Dentressangle et al. 2008, Velando and Torres 2014). Female food intake may also be under the indirect influence of her mate, if, for example, his attendance at the nest site frees her to spend more time foraging before and during egg laying. Male and Female Age acted additively on all traits except Breeding Date, where breeding delays in old age were accentuated for females paired with a young male and nearly disappeared for the partners of middle-aged and older males.

### Effects of the El Niño-Southern Oscillation on breeding traits

Nazca boobies’ responses to ENSO-related variability began early in the breeding cycle and showed a complex array of trait- and age-dependence. Anomalously warm sea surface temperatures (as during an El Niño) reduce primary productivity in the eastern equatorial Pacific Ocean (Fiedler et al. 1992, Fiedler 2002), but – contrary to predictions – larger Clutch Sizes and earlier Breeding Dates (for young birds only) accompanied positive SSTAs. Apparently beneficial effects of warm surface water on egg-laying traits preceded negative effects on Fledging Success (Anderson 1989, Tompkins and Anderson 2019) and juvenile survival (Champagnon et al. 2018). The ENSO’s influence on seabird populations is typically indirect, mediated by food availability (e.g., Schreiber and Schreiber 1984, Anderson 1989, Ancona et al. 2012). Nazca boobies eat mostly sardines (*Sardinops sagax*) and flying fish (Exoceotidae; Anderson 1989, Tompkins et al. 2017). Flying fish are poorly studied, but female (not male) Nazca boobies captured smaller individuals during the moderate 1986-87 El Niño (Anderson 1989), perhaps signaling negative effects on fish populations. Loss of sardines was implicated in the reproductive failure of Galápagos sea lions (*Zalophus wollebaeki*; Trillmich et al. 1985) and blue-footed boobies (Anderson 1989) during two El Niños from the 1980s, so that restricted sardine availability may be a general feature of El Niño in Galápagos. Thus, the two dominant taxa present in the Nazca booby diet each shows some indication of responding negatively to El Niño warm events. Poor food availability during the El Niño phase offers a potential explanation for low offspring production and juvenile survival, but not for large Clutch Size and early breeding in young birds (this study) and during the 1986-87 event (Anderson 1989). Nazca boobies participate in facilitated foraging with tuna (Spear et al. 2007, Johnston 2011), and extensive damage to the feet and legs of Nazca boobies is evidence of their interactions with predatory fish, dolphins, and/or sharks (Zavalaga et al. 2012). Tuna-dolphin-seabird foraging flocks are more common in the Galápagos Marine Reserve (133,000 sq km surrounding the Galápagos Islands) when SSTs are relatively warm (a seasonal effect; Johnston 2011), and we speculate that warming waters at the start of an El Niño event may enhance access to prey in advance of any negative effects on prey populations. Indeed, Nazca boobies were marginally more likely to regurgitate food items in Nov.-Jan. when sea surface temperature were relatively warm (β = 0.30 [95% CI: −0.04, 0.63]), but additional research is required to clarify ecological effects on biotic interactions in this system.

## ACKNOWLEDGEMENTS

We thank the Galapagos National Park Service for permission to work in the Park; the Charles Darwin Research Station, and TAME Airline, for logistical support; the National Geographic Society and Wake Forest University for research funding; and Nigel G. Yoccoz and three anonymous reviewers for comments which improved the manuscript. This material is based upon work supported primarily by the National Science Foundation under Grant Nos. DEB 93045679, DEB 9629539, DEB 98–06606, DEB 0235818, DEB 0842199, and DEB 1354473 to DJA. This publication is contribution #### of the Charles Darwin Foundation for the Galapagos Islands.

## REFERENCES

1. Ancona, S., I. Calixto-Albarrán, and H. Drummond. 2012. Effect of El Niño on the diet of a specialist seabird, *Sula nebouxii*, in the warm eastern tropical Pacific. Marine Ecology Progress Series 462:261–271.

2. Ancona, S., S. Sánchez-Colón, C. Rodríguez, and H. Drummond. 2011. El Niño in the warm tropics: local sea temperature predicts breeding parameters and growth of blue-footed boobies. Journal of Animal Ecology 80:799–808.

3. Anderson, D. J. 1989. Differential responses of boobies and other seabirds in the Galápagos to the 1986-87 El Niño-Southern Oscillation event. Marine Ecology Progress Series 52:209–16.

4. Anderson, D. J. 1990. Evolution of obligate siblicide in boobies. 1. A test of the insurance-egg hypothesis. The American Naturalist 135:334–350.

5. Anderson, D. J., and R. E. Ricklefs. 1987. Radio-tracking masked and blue-footed boobies (Sula spp.) in the Galápagos Islands. National Geographic Research 3:152–163.

6. Angelier, F., H. Weimerskirch, S. Dano, and O. Chastel. 2007. Age, experience and reproductive performance in a long-lived bird: a hormonal perspective. Behavioral Ecology and Sociobiology 61:611–621.

7. Apanius, V., M. Westbrock, and D. Anderson. 2008. Reproduction and immune homeostasis in a long-lived seabird, the Nazca booby (Sula granti). Ornithological Monographs 65:1–46.

8. Auld, J. R., and A. Charmantier. 2011. Life history of breeding partners alters age-related changes of reproductive traits in a natural population of blue tits. Oikos 120:1129–1138.

9. Auld, J. R., C. M. Perrins, and A. Charmantier. 2013. Who wears the pants in a mute swan pair? Deciphering the effects of male and female age and identity on breeding success. Journal of Animal Ecology 82:826–35.

10. Barber, R. T., and F. P. Chavez. 1983. Biological consequences of El Niño. Science 222:1203–1210.

11. Barbraud, C., and H. Weimerskirch. 2005. Environmental conditions and breeding experience affect costs of reproduction in blue petrels. Ecology 86:682–692.

12. Beamonte-Barrientos, R., A. Velando, H. Drummond, and R. Torres. 2010. Senescence of maternal effects: aging influences egg quality and rearing capacities of a long-lived bird. The American Naturalist 175:469–80.

13. Becker, P. H., and S. J. Bradley. 2007. The role of intrinsic factors for the recruitment process in long-lived birds. Journal of Ornithology 148:377–384.

14. Berman, M., J.-M. Gaillard, and H. Weimerskirch. 2009. Contrasted patterns of age-specific reproduction in long-lived seabirds. Proceedings of the Royal Society B: Biological Sciences 276:375–382.

15. Bjørkvoll, E., A. M. Lee, V. Grøtan, B. E. Sæther, A. Stien, S. Engen, S. Albon, L. E. Loe, and B. B. Hansen. 2016. Demographic buffering of life histories? Implications of the choice of measurement scale. Ecology 97:40–47.

16. Bonenfant, C., J.-M. Gaillard, F. Klein, and A. Loison. 2002. Sex- and age-dependent effects of population density on life history traits of red deer. Ecography 25:446–458.

17. Brommer, J. E., and K. Rattiste. 2008. “Hidden” reproductive conflict between mates in a wild bird population. Evolution 62:2326–33.

18. Bunce, A., S. J. Ward, and F. I. Norman. 2005. Are age-related variations in breeding performance greatest when food availability is limited? Journal of Zoology 266:163–169.

19. Cai, W., et al. 2014. Increasing frequency of extreme El Niño events due to greenhouse warming. Nature Climate Change 4:111–116.

20. Cam, E., W. A. Link, E. G. Cooch, J. Monnat, and E. Danchin. 2002. Individual covariation in life-history traits: seeing the trees despite the forest. The American Naturalist 159:96–105.

21. Carpenter, B., A. Gelman, M. D. Hoffman, D. Lee, B. Goodrich, M. Betancourt, M. Brubaker, J. Guo, P. Li, and A. Riddell. 2017. Stan: A probabilistic programming language. Journal of Statistical Software 76:1–32.

22. Catry, P., and R. W. Furness. 1999. The influence of adult age on territorial attendance by breeding Great Skuas Catharacta skua: an experimental study. Journal of Avian Biology 30:399–406.

23. Champagnon, J. C., J. D. Lebreton, H. Drummond, and D. J. Anderson. 2018. Pacific decadal and El Niño oscillations shape survival of a seabird. Ecology 99:1063–1072.

24. Clifford, L. D., and D. J. Anderson. 2001. Food limitation explains most clutch size variation in the Nazca booby. Journal of Animal Ecology 70:539–545.

25. Clifford, L. D., and D. J. Anderson. 2002. Clutch size variation in the Nazca booby: a test of the egg quality hypothesis. Behavioral Ecology 13:274.

26. Clutton-Brock, T. H. 1984. Reproductive effort and terminal investment in iteroparous animals. The American Naturalist 123:212–229.

27. Clutton-Brock, T. H., S. D. Albon, and F. E. Guinness. 1987. Interactions Between Population Density and Maternal Characteristics Affecting Fecundity and Juvenile Survival in Red Deer. Journal of Animal Ecology 56:857–871.

28. Coulson, T., E. Catchpole, S. D. Albon, B. Morgan, J. Pemberton, T. H. Clutton-Brock, M. J. Crawley, and B. T. Grenfell. 2001. Age, sex, density, winter weather, and population crashes in Soay sheep. Science 292:1528–1531.

29. Cubaynes, S., P. F. Doherty, E. A. Schreiber, and O. Gimenez. 2011. To breed or not to breed: a seabird’s response to extreme climatic events. Biology Letters 7:303–6.

30. Curio, E. 1983. Why do young birds reproduce less well? Ibis 125:400–404.

31. Dentressangle, F., L. Boeck, and R. Torres. 2008. Maternal investment in eggs is affected by male feet colour and breeding conditions in the blue-footed booby, Sula nebouxii. Behavioral Ecology and Sociobiology 62:1899–1908.

32. Devney, C. A., M. Short, and B. C. Congdon. 2009. Sensitivity of tropical seabirds to El Niño precursors. Ecology 90:1175–1183.

33. Doherty, P. F., E. A. Schreiber, J. D. Nichols, J. E. Hines, W. A. Link, G. A. Schenk, and R. W. Schreiber. 2004. Testing life history predictions in a long-lived seabird: a population matrix approach with improved parameter estimation. Oikos 105:606–618.

34. Drent, A. R. H., and S. Daan. 1980. The prudent parent: energetic adjustments in avian b. Ardea 68:225–252.

35. Elliott, K. H., K. M. O’Reilly, S. A. Hatch, A. J. Gaston, J. F. Hare, and W. G. Anderson. 2014. The prudent parent meets old age: a high stress response in very old seabirds supports the terminal restraint hypothesis. Hormones and Behavior 66:828–837.

36. Erikstad, K. E., P. Fauchald, T. Tveraa, and H. Steen. 1998. On the cost of reproduction in long-lived birds: the influence of environmental variability. Ecology 79:1781–1788.

37. Feldman, G. C., D. Clark, and D. Halpern. 1984. Satellite color observations of the phytoplankton distribution in the eastern equatorial Pacific during the 1982-1983 El Niño. Science 226:1069–1071.

38. Fiedler, P. 2002. Environmental change in the eastern tropical Pacific Ocean: review of ENSO and decadal variability. Marine Ecology Progress Series 244:265–283.

39. Fiedler, P. C., F. P. Chavez, D. W. Behringer, and S. B. Reilly. 1992. Physical and biological effects of Los Niños in the eastern tropical Pacific, 1986-1989. Deep Sea Research 39:199–219.

40. Froy, H., R. Phillips, A. G. Wood, D. H. Nussey, and S. Lewis. 2013. Age-related variation in reproductive traits in the wandering albatross: evidence for terminal improvement following senescence. Ecology letters 16:642–9.

41. Gaillard, J., M. Festa-Bianchet, N. G. Yoccoz, A. Loison, and C. Toigo. 2000. Temporal variation in fitness components and population dynamics of large herbivores. Annual Review of Ecology and Systematics 31:367–393.

42. Goodrich, B., J. Gabry, I. Ali & S. Brilleman. (2016). rstanarm: Bayesian applied regression modeling via Stan. R package version 2.13.1. http://mc-stan.org/.

43. Hernández, N., M. Genovart, J. M. Igual, and D. Oro. 2015. The influence of environmental conditions on the age pattern in breeding performance in a transequatorial migratory seabird. Frontiers in Ecology and Evolution 3:69.

44. Hooten, M., and N. Hobbs. 2015. A guide to Bayesian model selection for ecologists. Ecological Monographs 85:3–28.

45. Huyvaert, K. P., and D. J. Anderson. 2004. Limited dispersal by Nazca boobies Sula granti. Journal of Avian Biology 35:46–53.

46. Jahncke, J., and E. Goya. 2000. Responses of three booby species to El Niño 1997-1998. The International Journal of Waterbird Biology 23:102–108.

47. Johnston, M. 2011. Tuna-dolphin-bird feeding assemblages in the Galapagos Marine Reserve and their response to the physical characteristics of the upper water column. Master’s Thesis, Texas A and M University.

48. Kim, S. Y., A. Velando, R. Torres, and H. Drummond. 2011. Effects of recruiting age on senescence, lifespan and lifetime reproductive success in a long-lived seabird. Oecologia 166:615–626.

49. Laaksonen, T., E. Korpimaki, and H. Hakkarainen. 2002. Interactive effects of parental age and environmental variation on the breeding performance of Tengmalm’s owls. Journal of Animal Ecology 71:23–31.

50. Lecomte, V. J., et al. 2010. Patterns of aging in the long-lived wandering albatross. Proceedings of the National Academy of Sciences (USA) 107: 6370–5.

51. Lemaître, J.-F., and J. M. Gaillard. 2017. Reproductive senescence: new perspectives in the wild. Biological Reviews 92:2182–2199.

52. Maness, T. J., and D. J. Anderson. 2007. Serial monogamy and sex ratio bias in Nazca boobies. Proceedings of the Royal Society B: Biological Sciences 274:2047–54.

53. Maness, T. J., and D. J. Anderson. 2008. Mate rotation by female choice and coercive divorce in Nazca boobies, *Sula granti*. Animal Behaviour 76:1267–1277.

54. Maness, T. J., and D. J. Anderson. 2013. Predictors of juvenile survival in birds. Ornithological Monographs 2013:1–55.

55. McNamara, J. M., A. I. Houston, Z. Barta, A. Scheuerlein, and L. Fromhage. 2009. Deterioration, death and the evolution of reproductive restraint in late life. Proceedings of the Royal Society B: Biological Sciences 276:4061–4066.

56. McPhaden, M. J., S. E. Zebiak, and M. H. Glantz. 2006. ENSO as an integrating concept in earth science. Science 314:1740–1745.

57. Nevoux, M., H. Weimerskirch, and C. Barbraud. 2007. Environmental variation and experience-related differences in the demography of the long-lived black-browed albatross. Journal of Animal Ecology 76:159–167.

58. Nur, N. 1984. Increased reproductive success with age in the California gull: due to increased effort or improvement of skill? Oikos 43:407–408.

59. Nussey, D. H., H. Froy, J.-F. Lemaître, J.-M. Gaillard, and S. N. Austad. 2013. Senescence in natural populations of animals: widespread evidence and its implications for bio-gerontology. Ageing Research Reviews 12:214–25.

60. Nussey, D. H., L. E. B. Kruuk, A. Morris, M. N. Clements, J. M. Pemberton, and T. H. Clutton-Brock. 2009. Inter- and intrasexual variation in aging patterns across reproductive traits in a wild red deer population. The American Naturalist 174:342–357.

61. Oro, D., N. Hernández, L. Jover, and M. Genovart. 2014. From recruitment to senescence: food shapes the age-dependent pattern of breeding performance in a long-lived bird. Ecology 95:446–457.

62. Oro, D., R. Torres, C. Rodríguez, and H. Drummond. 2010. Climatic influence on demographic parameters of a tropical seabird varies with age and sex. Ecology 91:1205–1214.

63. Pardo, D., C. Barbraud, M. Authier, and H. Weimerskirch. 2013. Evidence for an age-dependent influence of environmental variations on a long-lived seabird’s life-history traits. Ecology 94:208–220.

64. Pennington, J. T., K. L. Mahoney, V. S. Kuwahara, D. D. Kolber, R. Calienes, and F. P. Chavez. 2006. Primary production in the eastern tropical Pacific: a review. Progress in Oceanography 69:285–317.

65. van de Pol, M., and S. Verhulst. 2006. Age-dependent traits: a new statistical model to separate within- and between-individual effects. The American Naturalist 167:766–73.

66. R Core Team (2020). R: A language and environment for statistical computing. R Foundation for Statistical Computing, Vienna, Austria.

67. Ratcliffe, N., R. W. Furness, and K. C. Hamer. 1998. The interactive effects of age and food supply on the breeding ecology of great skua. Journal of Animal Ecology 67:853–862.

68. Schaffer, W. M. 1974. Optimal reproductive effort in fluctuating environments. The American Naturalist 108:783–790.

69. Schreiber, R. W., and R. A. Schreiber. 1984. Central Pacific seabirds and the El Niño Southern Oscillation: 1982 to 1983 perspectives. Science 222:713–716.

70. Spear, L. B., D. G. Ainley, W. A. Walker, C. D. Marti, and C. Valencia. 2007. Foraging Dynamics of Seabirds in the Eastern Tropical Pacific Ocean. Studies in Avian Biology 35:1–99.

71. Spear, L. B., L. T. Ballance, and D. G. Ainley. 2001. Response of seabirds to thermal boundries in the tropical Pacific: the thremocline verus the Equatorial front. Marine Ecology Progress Series 219:275–289.

72. Stearns, S. C. 1992. The Evolution of Life Histories. Oxford University Press, London.

73. Sydeman, W. J., J. F. Penniman, T. M. Penniman, P. Pyle, and D. G. Ainley. 1991. Breeding performance in the western gull: effects of parental age, timing of breeding and year in relation to food availability. Journal of Animal Ecology 60:135–149.

74. Tavecchia, G., T. Coulson, B. J. T. Morgan, J. M. Pemberton, J. C. Pilkington, F. M. D. Gulland, and T. H. Clutton-Brock. 2005. Predictors of reproductive cost in female Soay sheep. Journal of Animal Ecology 74:201–213.

75. Teplitsky, C., J. A. Mills, J. W. Yarrall, and J. Merilä. 2010. Indirect genetic effects in a sex-limited trait: the case of breeding time in red-billed gulls. Journal of Evolutionary Biology 23:935–44.

76. Tompkins, E. M., and D. J. Anderson. 2019. Sex-specific patterns of reproductive senescence in Nazca boobies linked to mating system. Journal of Animal Ecology 00:1–15.

77. Tompkins, E. M., H. M. Townsend, and D. J. Anderson. 2017. Decadal-scale variation in diet forecasts persistently poor breeding under ocean warming in a tropical seabird. PLoS ONE 12:e0182545.

78. Townsend, H. M., and D. J. Anderson. 2007a. Production of insurance eggs in Nazca boobies: costs, benefits, and variable parental quality. Behavioral Ecology 18:841–848.

79. Townsend, H. M., and D. J. Anderson. 2007b. Assessment of costs of reproduction in a pelagic seabird using multistate mark-recapture models. Evolution 61:1956–68.

80. Trenberth, K. E. 1997. The definition of El Niño. Bulletin of the American Meteorological Society 78:2771–2777.

81. Trillmich, F., D. Limberger, and S. Url. 1985. Drastic effects of El Niño on Galapagos pinnipeds. Oecologia 67:19–22.

82. Vehtari, A., A. Gelman, and J. Gabry. 2016. loo: Efficient leave-one-out cross-validation and WAIC for Bayesian models.

83. Velando, A., H. Drummond, and R. Torres. 2006. Senescent birds redouble reproductive effort when ill: confirmation of the terminal investment hypothesis. Proceedings of the Royal Society B: Biological Sciences 273:1443–1448.

84. Velando, A., and R. Torres. 2014. Enhanced male coloration after immune challenge increases reproductive potential. Journal of Evolutionary Biology 27:1582–1589.

85. Vieyra, L., E. Velarde, and E. Ezcurra. 2009. Effects of parental age and food availability on the reproductive success of Heermann’s gulls in the Gulf of California. Ecology 90:1084–1094.

86. Wang, C., and P. C. Fiedler. 2006. ENSO variability and the eastern tropical Pacific: a review. Progress in Oceanography 69:239–266.

87. Williams, G. 1966. Natural selection, the costs of reproduction, and a refinement of Lack’s principle. The American Naturalist 100:687–690.

88. Weimerskirch, H. 1992. Reproductive Effort in Long-Lived Birds : Age-Specific Patterns of Condition, Reproduction and Survival in the Wandering Albatross. Oikos 64:464–473.

89. Wood, S. N. 2006. Generalized additive models: an introduction with R. CRC Press, Boca Raton, FL, USA.

90. Yeh, S. W., J. S. Kug, B. Dewitte, M. H. Kwon, B. P. Kirtman, and F. F. Jin. 2009. El Niño in a changing climate. Nature 461:511–514.

91. Zavalaga, C. B., S. D. Emslie, F. A. Estela, M. S. Müller, G. Dell’Omo, and D. J. Anderson. 2012. Overnight foraging trips by chick-rearing Nazca Boobies *Sula granti* and the risk of attack by predatory fish. Ibis 154:61–73.

