## Appendix S1 for "Breeding responses to environmental variation are age- and trait-dependent in female Nazca boobies"

### APPENDIX S1 – Methods: data selection and adjustments to cope with missing information

In most years, some Nazca booby clutches (primarily renests) were laid in February or March, months after the seasonal peak in laying (November/December) had subsided. These late clutches were typically abandoned quickly and did not produce surviving offspring (unpublished data). Late clutches were excluded from analyses because they were not monitored to the same standard in all years. Three-egg clutches (< 0.5% of clutches) were excluded from clutch size analyses, and rare cases when eggs or nestlings were adopted into a clutch (< 2% of clutches) were excluded from Fledging Success analyses.

In many seasons, some clutches were established before colony monitors arrive. For these clutches, Breeding Dates were estimated from the laying date of the second-laid egg or from hatching date(s) using the average laying asynchrony (5 days, length of time between laying of the first and second eggs in two-egg clutches; Anderson 1989) and/or the average incubation period (43 days; unpublished data). For clutches laid before monitor arrival and lacking hatching dates, Breeding Date was assigned as the median laying date for clutches 1) laid before monitors arrived and 2) hatching. Across all years, 2.4% of records have Breeding Date assigned this way. The 1998 and 1999 seasons (a strong La Niña event) are a special case: following the extreme El Niño in 1997-98, breeding was unusually early at the Punta Cevallos colony and hatching success was low, so that 33% and 20% (respectively) of Breeding Dates are missing. All Breeding Date analyses exclude 1998 and 1999. Breeding Probability analyses also excluded these two years (1998 and 1999) because some clutches may have been laid, and failed, before monitors observed them, leading to errors in the assignment of breeder/non-breeder status for banded females. Breeding Probability analyses also excluded 2017 because nest initiations in Subcolony 3 were not monitored comprehensively in that year (Subcolony 3 is one of three monitored subcolonies; Huyvaert and Anderson 2004), so that breeder/non-breeder status could not be assigned with certainty for all living individuals.

When two-egg clutches were established before colony monitors arrived, order of laying was not available to distinguish A-eggs from B-eggs. Hatching dates established egg identities for some affected clutches. For others, the larger egg was assumed to be the first laid because A-egg volume ≥ B-egg volume in 90% of clutches with known laying order. For the rest, B-egg volumes and A-egg volumes were similar so that the error introduced by a misidentification is small: only 10% of cases where the B-egg volume exceeds the A-egg volume have a B-egg volume > 10% larger).

In some breeding seasons included in this study (2010-2017), not all young-of-the-year reached independence before colony monitoring ended. For these seasons, Fledging Success (the probability of an offspring reaching independence, given that a nest was initiated) was adjusted based on the recovery of banded offspring carcasses during the following breeding season (5-6 months later). Using this approach, some offspring may die but be classified as “reaching independence” if carcass recoveries are not complete. We evaluated this possibility using data from two seasons. In the 2010-11 breeding season, dead chick carcasses were located a few weeks after monitoring ceased (in mid-May) and used as a starting sample from which to calculate the proportion of carcasses that were re-located in the following season. Twenty-seven banded offspring carcasses were present in May 2011, and 20 of these were later recovered (recovery rate = 74%). In 2015, an unusually high number of offspring died late in the breeding season, all these young-of-the-year carcasses were left in place and were used to estimate the rate of band recovery the following year. Two hundred seventeen carcasses were left in place, and 122 of these were recovered (recovery rate = 56%). The recovery rate in 2015 may have been lower than that from 2010 for three reasons: (1) because chicks were left in place from March-May in 2015, some of the carcasses had to persist for 1-2 months longer in 2015 vs. in 2010 before the detection period the following fall, (2) the 2015 carcasses included younger, smaller, chicks whose carcasses may be more prone to disarticulate and/or disperse, and (3) the 2010-11 carcasses were located once after death (in May) to enter the sample, which may have biased the starting sample toward carcasses that were relatively easy to find. The rates of recovery estimated in these two seasons (56% recovery in 2015, 74% recovery in 2010) were used to look at the effect of this error on annual rates of Fledging Success for all years 2010-2017 (Table S1). To do so, we calculated the number of offspring that died, but that will be misclassified as reaching independence due to not finding their carcass after death. We then calculated the effect of the predicted misclassifications on mean Fledging Success for each season (Table S1).

While detection rates for dead chick carcasses are far below 1, relatively few offspring die after the monitoring period ends (Table S1), such that the effect of imperfect carcass detection on estimated Fledging Success is negligible. For 56% recovery, Fledging Success will be overestimated by 0.2-2.4%, and for 74% recovery, Fledging Success will be overestimated by 0.1-1.2%. In Table S1, the number of chick deaths detected (the first row) includes all banded chicks (not just those from known-age, banded mothers, included in this study).

**Table S1**. Young-of-the-year deaths detected only from band recoveries in a later season and the implications of low (56%) or high (74%) rates of band recovery for Fledging Success.

|  | **Breeding Season** | | | | | | | |
| --- | --- | --- | --- | --- | --- | --- | --- | --- |
|  | **2010** | **2011** | **2012** | **2013** | **2014** | **2015** | **2016** | **2017** |
| # of chick deaths detected by locating a banded carcass in a later season | 44 | 20 | 7 | 70 | 39 | 50 | 23 | 27 |
| # of offspring assumed to reach independence ("fledged") | 558 | 1,273 | 589 | 889 | 1,443 | 960 | 389 | 1,171 |
| The proportion of offspring misclassified as fledged under a 0.56 detection frequency | 0.055 | 0.012 | 0.008 | 0.057 | 0.020 | 0.032 | 0.039 | 0.017 |
| The proportion of offspring misclassified as fledged under a 0.74 detection frequency | 0.028 | 0.005 | 0.004 | 0.028 | 0.009 | 0.018 | 0.021 | 0.008 |
| The degree of overestimation in Fledging Success (56% detection frequency) | 0.015 | 0.006 | 0.002 | 0.024 | 0.013 | 0.013 | 0.008 | 0.013 |
| The degree of overestimation in Fledging Success (74% detection frequency) | 0.008 | 0.003 | 0.001 | 0.012 | 0.006 | 0.008 | 0.004 | 0.006 |

*References*

Anderson, D. J. 1989. The role of hatching asynchrony in siblicidal brood reduction of two booby species. Behavioral Ecology and Sociobiology 25:363-368.
