## Appendix S2 for "Breeding responses to environmental variation are age- and trait-dependent in female Nazca boobies"

### APPENDIX S2 – Time series of environmental predictor variables

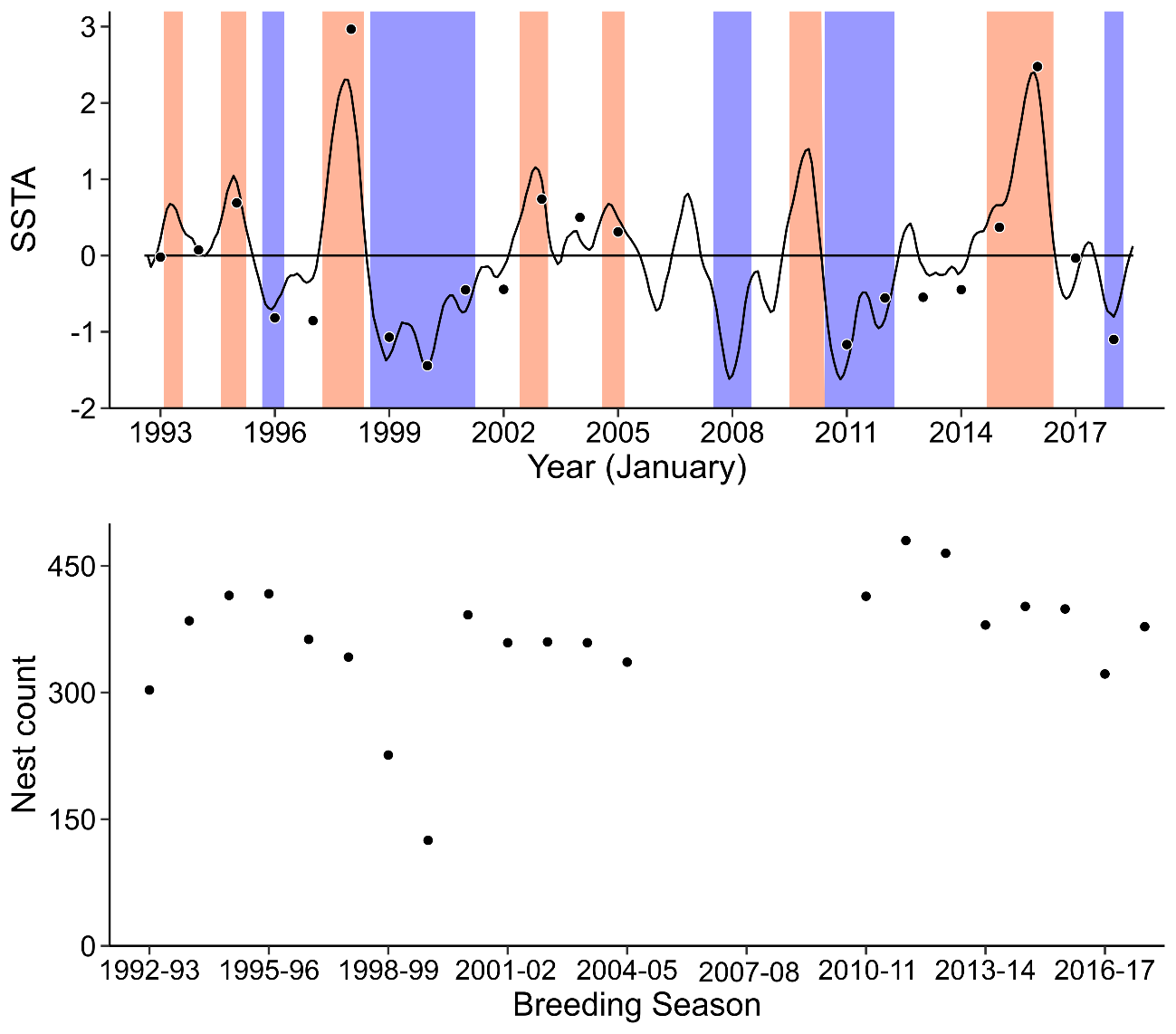

**Figure S1.** Time series for two continuous measures of the quality of the Nazca booby breeding environment. Panel (a) shows average December-February monthly SSTA from the Niño 3 region (black dots) overlaid on the categorical occurrence of El Niño/La Niña (shaded red/blue, respectively) following the definition of Trenberth (1997) using monthly SSTA from the Niño 3.4 region (solid line). Nazca booby breeding seasons span parts of two calendar years. SSTA values in (a) are labelled by calendar year (in January) and occur during the breeding season starting the previous fall: e.g., the SSTA value over January 1993 falls within the 1992-93 breeding season. Panel (b) shows the annual number of nest initiations in a comprehensively monitored colony subsection, with re-nesting attempts excluded.

*References*

Trenberth, K. E. 1997. The definition of El Niño. Bulletin of the American Meteorological Society 78:2771–2777.
