## Appendix S3 for "Breeding responses to environmental variation are age- and trait-dependent in female Nazca boobies"

### APPENDIX S3 – Methods: regurgitation data

In some breeding seasons we have collected monthly diet samples by regurgitation from a small number of Nazca booby adults (usually 20) during egg-laying (November-January). Diet samples were collected from (randomly selected) adults returning to the colony from a foraging trip in the late afternoon (1700-1900 h). We record whether each bird regurgitates one or more food items and, for successful regurgitations, the number and identity of food items (to species if possible) and a sample mass. During November, December, and January, many birds did not regurgitate (67%), and we used the probability of regurgitating one or more food items, given that a bird is sampled, as an index of food availability.

To support a relationship between SSTA and resource availability, we fit a binomial GLMM to diet data, predicting the probability of regurgitating by SSTA and SSTA^2^. Data were available from 1992-1994, 1999-2004, and 2011-2017 only (969 sampled birds), and not all breeding seasons contributed November and/or December samples. Additional predictors of regurgitation probability included Fish Phase (Tompkins & Anderson 2017) and the date of sampling relative to the season-specific mean breeding date (“Relative Sampling Date”). Sampling session (month nested within year) was included as a random intercept. We sampled roughly equal numbers of males and females in each session and we did not include Sex as a predictor of regurgitation probability (doing so would exclude a few records missing data on sex).
