## Appendix S4 for "Breeding responses to environmental variation are age- and trait-dependent in female Nazca boobies"

### APPENDIX S4 – Sample sizes by response variable and analysis

**Table S1.** Sample sizes organized by analysis and response variable.

| *Age plus environment* | | | | | |
| --- | --- | --- | --- | --- | --- |
| **Sample size** |  | **Breeding Date** | **Clutch Size** | **A-egg Volume** |  |
| Breeding seasons |  | 20 | 20 | 20 |  |
| Females |  | 2,659 | 2,658 | 2,648 |  |
| Records |  | 14,860 | 14,949 | 14,383 |  |
| *Selective disappearance* | | | | | |
| **Sample size** |  | **Breeding Date** | **Clutch Size** | **A-egg Volume** |  |
| Breeding seasons |  | 20 | 20 | 20 |  |
| Females |  | 473 | 474 | 470 |  |
| Records |  | 3,134 | 3,285 | 3,011 |  |
| *Male influence* | | | | | |
| **Sample size** |  | **Breeding Date** | **Clutch Size** | **A-egg Volume** | **Fledging Success** |
| Breeding seasons |  | 20 | 20 | 20 | 20 |
| Females |  | 1,965 | 1,962 | 1,952 | 1,949 |
| Males |  | 2,248 | 2,248 | 2,232 | 2,226 |
| Records |  | 7,947 | 7,929 | 7,801 | 7,662 |
| *Age by environment interactions, ages 3-12* | | | | | |
| **Sample size** | **Breeding Probability** | **Breeding Date** | **Clutch Size** | **A-egg Volume** | **Fledging Success** |
| Breeding seasons | 17 | 18 | 20 | 20 | 20 |
| Females | 2,918 | 2,654 | 2,653 | 2,643 | 2,641 |
| Records | 14,558 | 11,264 | 11,337 | 10,902 | 10,676 |
| *Age by environment interactions, ages 11-20* | | | | | |
| **Sample size** | **Breeding Probability** | **Breeding Date** | **Clutch Size** | **A-egg Volume** | **Fledging Success** |
| Breeding seasons | 7 | 8 | 8 | 8 | 8 |
| Females | 1,437 | 1,518 | 1,517 | 1,508 | 1,505 |
| Records | 4,664 | 4,926 | 4,898 | 4,798 | 4,699 |
