## Appendix S5 for "Breeding responses to environmental variation are age- and trait-dependent in female Nazca boobies"

APPENDIX S5 – Methods: details of threshold age function parameterization

Two-threshold models for each response variable used the following parameterization of the linear predictor for individual *i* at age *j* (shown here for Clutch Size):

logit(ClutchSize*_ij_*) = α + β_1_Age*_ij_* + β_2_(Age*_ij_* – T_1_)_+_ + β_3_(Age*_ij_* – T_2_)_+_ + β_4_x_4_…, + β*_k_*x*_k_ + u_i_* + *u_yr_*

Where threshold ages T_1_ (6, 7, 8, or 9 years old) and T_2_ (13, 14, 15, 16, or 17 years old) were set for each candidate model and (Age*_ij_* – T_1_)_+_ represents the product of (Age*_ij_* – T_1_) and a logical function: step(Age*_ij_* ≥ T_1_). ‘step(Age*_ij_* ≥ T_1_)’ equals 1 when Age*_ij_* ≥ T_t_, and 0 otherwise. Coefficient β_1_ represents the slope of each response variable on age for the early life period (ages ≥ T_1_), β_2_ represents the change in slope after the first threshold age, and β_3_ represents the change in slope after the second threshold age. The logical function toggles the β_2_ term *off* for ages ≤ T_1_ and *on* for ages > than T_1_. In this example, the slope of Clutch Size on age before T_1_ is β_1_, the slope of Clutch Size on age between T_1_ and T_2_ is β_1_ + β_2,_ and the slope of Clutch Size on age after T_2_ is β_1+_ β_2+_ β_3_. Additional predictors (β_4_x_4_…, + β*_k_*x*_k_*) were included as described in the main text. Lastly, u*_i_* and u*_yr_* are individual- and year-specific group-level effects.
