## Appendix S6 for "Breeding responses to environmental variation are age- and trait-dependent in female Nazca boobies"

### APPENDIX S6 – Interpretation of Nest Count

We assumed that Nest Count, the number of first clutch initiations in a comprehensively monitored subsection of the colony, positively covaried with the quality of the breeding environment. However, the number of competitors may increase with the number of breeding pairs, introducing negative effects on prey availability through enhanced intraspecific competition (Lewis et al. 2001). Negative effects of competitive interactions may be greatest for young (Oro et al. 2014) or old individuals (Lecomte et al. 2010). Relatively high levels of intraspecific competition should result in greater foraging trip distance and longer trip duration (Lewis et al. 2001). Trip distance and trip duration are highly correlated in masked boobies (*Sula dactylatra*; Weimerskirch et al. 2008) and we used 5 years of detailed data on the nest attendance of incubating Nazca boobies to approximate foraging trip duration and test the idea that Nest Count positively associates with the length of time females are absent from the nest (presumably on a foraging trip, variable “Absence Duration”). If density dependent processes outweigh the influence of food availability on Nest Count, we expect Absence Duration to increase with the number of breeding pairs, particularly for young and old individuals.

In breeding seasons 2010-2014 the sex of the incubating parent was recorded daily for all nests in the Study Area, a comprehensively monitored subsection of the colony (Huyvaert and Anderson 2004, Apanius et al. 2008). Male and female pair members take turns incubating their clutch. A female was considered absent from the nest (presumably foraging) when the male was recorded incubating; the length of her absence (in days) was defined as the number of consecutive days the male attended the nest. Absence Duration was defined as the mean length of each female’s absences across all records for the nest (excluding the last incubation stint before clutch failure, clutch hatching, or the end of the monitoring period). Daily nest checks spanned 4 November-9 January in the 2010 season, 5 October-9 January in the 2011 season, 29 September-7 January in the 2012 season, 20 October-5 January in the 2013 season, and 24 October-11 January in the 2014 season. Only Absence Durations for known-age females performing > 3 incubation stints were included in the data. Nest Count varied from 380 clutch initiations to 480 clutch initiations across the five seasons included in the data.

The influence of Nest Count on Absence Duration was modelled using a linear mixed model with Breeding Season and Female Identity fit as random intercepts. Nest Count (standardized to a mean of zero and unit s.d.) was fit as a linear predictor. Models also included Female Age (fit as a categorical variable because our interest was not in establishing age effects on Absence Duration) and an interaction between Female Age and Nest Count.

Nest count did not influence Absence Duration (β = -0.20 [95% BCI: -0.66, 0.30], from an LMM excluding a Female Age x Nest Count interaction) but it is worth noting that the relationship between these two variables, while not statistically supported, suggests that Absence Durations get shorter, not longer (as expected under density-dependence) with increasing Nest Count. Old females (≥ 16 years of age) and young females (≤ 7 years of age) had longer Absence Durations than middle-aged birds. Relationships between Nest Count and Absence Durations did not vary by age category (Table S1). Absence Durations were measured on a coarse scale in this analysis and the data were limited to only five years, nevertheless, these preliminary results give no support to the idea that negative effects of increased density increase with Nest Count.

**Table S1**. Coefficient estimates from the most-general LMM explaining variation in Absence Duration by Nest Count and categorical female age. Coefficients distinct from zero are in bold.

| Coefficients | Mean [95% BCI] |
| --- | --- |
| **Intercept** | **2.51 [2.04, 2.99]** |
| **Categorical F Age (old)** | **0.19 [0.06, 0.33]** |
| **Categorical F Age (young)** | **0.19 [0.09, 0.30]** |
| Nest Count | -0.21 [-0.69, 0.26] |
| F Age (old) x Nest Count | 0.13 [-0.01, 0.26] |
| F Age (young) x Nest Count | -0.03 [-0.14, 0.08] |
| Group-specific effects | Var. [95% BCI] |
| Female ID | 0.02 [0.00, 0.05] |
| Year | 0.27 [0.04, 1.26] |
| Residual | 0.68 [0.65, 0.72] |

*References*

Apanius, V., M. Westbrock, and D. Anderson. 2008. Reproduction and immune homeostasis in a long-lived seabird, the Nazca booby (*Sula granti*). Ornithological Monographs 65:1–46.

Huyvaert, K. P., and D. J. Anderson. 2004. Limited dispersal by Nazca boobies Sula granti. Journal of Avian Biology 35:46–53.

Lecomte, V. J., G. Sorci, S. Cornet, A. Jaeger, B. Faivre, E. Arnoux, M. Gaillard, C. Trouvé, D. Besson, O. Chastel, and H. Weimerskirch. 2010. Patterns of aging in the long-lived wandering albatross. Proceedings of the National Academy of Sciences of the United States of America 107:6370–5.

Lewis, S., T. N. Sherratt, K. C. Hamer, and S. Wanless. 2001. Evidence of intra-specific competition for food in a pelagic seabird. Nature 412:816–819.

Oro, D., N. Hernández, L. Jover, and M. Genovart. 2014. From recruitment to senescence: Food shapes the age-dependent pattern of breeding performance in a long-lived bird. Ecology 95:446–457.

Weimerskirch, H., M. Le Corre, and C. A. Bost. 2008. Foraging strategy of masked boobies from the largest colony in the world: Relationship to environmental conditions and fisheries. Marine Ecology Progress Series 362:291–302.
