## Appendix S7 for "Breeding responses to environmental variation are age- and trait-dependent in female Nazca boobies"

### APPENDIX S7 – Methods: priors, posterior estimation, and model fit

Program Stan (Carpenter et al. 2017) efficiently samples a model’s posterior probability distributions using an adaptive variant (the No U-Turn Sampler, or NUTS) of Hamiltonian Monte Carlo, an algorithm suited to complex models containing many parameters. We used four chains to sample the posterior distribution for each model, running 3,000 warm-up iterations and then a further 3,000 sampling iterations with a thinning interval of six. We retained 2,000 posterior samples per model, which were used to calculate marginal posterior means and credible intervals for parameters. Estimated sample size (accounting for the autocorrelation inherent to MCMC chains) was always greater than 1,500 and often equal to 2,000, justifying our choice of sampling procedure. For each model, chain convergence was diagnosed visually and using the potential scale reduction criteria of Gelman et al. (2013). Fitted models were further evaluated using posterior predictive checks (Gabry et al. 2017).

We set weakly regularizing priors on the fixed effect intercept and slope coefficients using a normal (µ = 0, σ = 10) prior for covariates fit to Breeding Date and A-egg Volume and a normal (µ = 0, σ = 2.5) prior for covariates fit to Breeding Probability, Clutch Size, and Fledging Success. For all response variables, we used the default “decomposition of covariance” prior from the rstanarm package (v. 2.13.1) on the random effects (Goodrich et al., 2016), which is equivalent to setting a unit exponential prior on the standard deviation for Breeding Season and Identity random intercepts. Posterior means and 95% CI estimates were not different under the stronger versus weaker priors (data not shown). We used the default unit exponential prior on the standard deviation of the smooth terms for fitted GAMMs.

*References*

Carpenter, B., A. Gelman, M. D. Hoffman, D. Lee, B. Goodrich, M. Betancourt, M. Brubaker, J. Guo, P. Li, and A. Riddell. 2017. Stan: A probabilistic programming language. Journal of Statistical Software 76:1–32.

Gabry, J., D. Simpson, A. Vehtari, M. Betancourt, and A. Gelman. 2017. Visualization in Bayesian workflow. arXiv:1709.01449.

Gelman, A., J. B. Carlin, H. S. Stern, D. B. Dunson, A. Vehtari, and D. B. Rubin. 2013. *Bayesian Data Analysis*, Third Edit. CRC Press, Boca Raton, FL.

Goodrich, B., J. Gabry, I. Ali & S. Brilleman. (2016). rstanarm: Bayesian applied regression modeling via Stan. R package version 2.13.1. <http://mc-stan.org/>.
