## Appendix S8 for "Breeding responses to environmental variation are age- and trait-dependent in female Nazca boobies"

### APPENDIX S8 – Results: model selection results and coefficient estimates by response variable and analysis stage

**Table S1**. Model selection results support female age effects on Breeding Date, Clutch Size, and A-egg Volume in Nazca boobies. Models were ranked by their out-of-sample prediction accuracy using the expected log predictive densities from 10-fold cross-validation (*elp̂d_10-fold CV_*). Models with higher (less negative) *elp̂d_10-fold CV_* have better explanatory ability. Δ*elp*̂*d_10-fold CV_* gives the distance, for each candidate model, from the best-supported GLMM. Model rankings compare alternative parameterizations of age effects, including a non-parametric smooth function (labelled “s(F Age)”, a GAMM) and single-threshold (“One T”) or two-threshold functions (“Two T”). Any model whose performance overlaps that of the best-supported GLMM is in bold. Female Identity and Breeding Season random effects and additional fixed-effect predictors were held constant across all candidate models (Methods)..

**Table S1**.

|  | Breeding Date | |  | A-egg Volume | |  | Clutch Size | |
| --- | --- | --- | --- | --- | --- | --- | --- | --- |
| **Age Model** | *Δelp̂d_10-fold CV_* (SE) | *elp̂d_10-fold CV_* (SE) |  | *Δelp̂d_10-fold CV_* (SE) | *elp̂d_10-fold CV_* (SE) |  | *Δelp̂d_10-fold CV_* (SE) | *elp̂d_10-fold CV_* (SE) |
| Age | -1,018.9 (49.9) | -16,419.6 (102.0) |  | -426.4 (51.5) | -14,480.9 (189.4) |  | -117.9 (16.3) | -8,735.3 (52.1) |
| Age + Age^2^ | -208.4 (30.4) | -15,609.1 (102.9) |  | -164.5 (40.9) | -14,219.0 (200.1) |  | -30.5 (9.9) | -8,647.9 (53.2) |
| No Age | -1,288.8 (57.5) | -16,689.5 (102.9) |  | -456.4 (46.7) | -14,510.9 (191.8) |  | -120.5 (16.4) | -8,737.9 (52.1) |
| One T (6) | -338.2 (32.9) | -15,738.9 (102.6) |  | -279.1 (43.9) | -14,333.6 (199.4) |  | -28.4 (8.7) | -8,645.8 (53.2) |
| One T (7) | -161.6 (25.3) | -15,562.3 (103.0) |  | -100.8 (18.6) | -14,155.3 (187.0) |  | -36.9 (9.0) | -8,654.3 (53.2) |
| One T (8) | -101.4 (26.5) | -15,502.1 (102.3) |  | -117.2 (42.1) | -14,171.7 (191.7) |  | -45.2 (9.9) | -8,662.6 (53.1) |
| One T (9) | -111.8 (29.1) | -15,512.4 (101.8) |  | -141.3 (52.3) | -14,195.8 (202.8) |  | -36.3 (10.2) | -8,653.8 (53.0) |
| s(F Age) | 61.6 (21.5) | -15,339.1 (102.9) |  | -48.1 (39.7) | -14,102.6 (201.1) |  | 4.4 (6.7) | -8,613.0 (53.3) |
| Two T (6, 13) | -84.3 (24.8) | -15,484.9 (102.2) |  | -127.3 (40.6) | -14,181.8 (202.8) |  | **0.0 (0.0)** | **-8,617.4 (53.5)** |
| Two T (6, 14) | -79.1 (24.9) | -15,479.8 (102.3) |  | -170.1 (45.3) | -14,224.6 (204.4) |  | **-5.9 (5.6)** | **-8,623.3 (53.5)** |
| Two T (6, 15) | -153.8 (25.1) | -15,554.4 (103.5) |  | -174.8 (54.2) | -14,229.2 (203.9) |  | **-9.3 (6.0)** | **-8,626.7 (53.5)** |
| Two T (6, 16) | -130.3 (26.5) | -15,531.0 (102.5) |  | -117.4 (53.9) | -14,171.9 (192.3) |  | **-11.9 (6.5)** | **-8,629.3 (53.5)** |
| Two T (6, 17) | -162 (27.3) | -15,562.7 (103.5) |  | -186.6 (47.2) | -14,241.0 (199.3) |  | **-5.1 (6.9)** | **-8,622.5 (53.4)** |
| Two T (7, 13) | **-7.5 (18.6)** | **-15,408.2 (103.1)** |  | **0.0 (0.0)** | **-14,054.5 (192.1)** |  | -15.1 (7.1) | -8,632.5 (53.2) |
| Two T (7, 14) | **-25.2 (18.4)** | **-15,425.9 (103.1)** |  | **-82.9 (43.9)** | **-14,137.4 (203.6)** |  | -19.3 (7.2) | -8,636.7 (53.4) |
| Two T (7, 15) | **0.0 (0.0)** | **-15,400.7 (103.3)** |  | -105.0 (42.9) | -14,159.5 (205.7) |  | -23.9 (7.4) | -8,641.3 (53.4) |
| Two T (7, 16) | **-16.3 (18.7)** | **-15,417.0 (103.2)** |  | **-75.8 (52.6)** | **-14,130.2 (195.3)** |  | -25.9 (7.6) | -8,643.3 (53.3) |
| Two T (7, 17) | **-15.1 (18.6)** | **-15,415.8 (103.2)** |  | **-77.6 (43.2)** | **-14,132.1 (203.0)** |  | **-13.3 (7.9)** | **-8,630.8 (53.2)** |
| Two T (8, 13) | **-48.1 (24.0)** | **-15,448.8 (102.3)** |  | **-46.9 (39.8)** | **-14,101.3 (193.2)** |  | -23.7 (8.7) | -8,641.1 (53.2) |
| Two T (8, 14) | **-12.5 (23.6)** | **-15,413.1 (102.3)** |  | **-57.4 (41.5)** | **-14,111.9 (202.6)** |  | -28.7 (8.8) | -8,646.1 (53.2) |
| Two T (8, 15) | **-35.6 (23.5)** | **-15,436.2 (102.2)** |  | -115.7 (43.3) | -14,170.2 (205.6) |  | -19.8 (8.8) | -8,637.2 (53.2) |
| Two T (8, 16) | **-25.3 (23.6)** | **-15,426.0 (102.8)** |  | **-72.6 (50.1)** | **-14,127.1 (197.7)** |  | -18.8 (8.9) | -8,636.2 (53.1) |
| Two T (8, 17) | **-2.7 (23.7)** | **-15,403.4 (102.8)** |  | **-62.5 (37.2)** | **-14,116.9 (198.6)** |  | -35.7 (9.2) | -8,653.1 (53.3) |
| Two T (9, 13) | -96.6 (28.6) | -15,497.3 (102.3) |  | -106.4 (54.7) | -14,160.9 (205.4) |  | -36.3 (9.6) | -8,653.7 (53.1) |
| Two T (9, 14) | -90.0 (28.2) | -15,490.7 (102.4) |  | -97.4 (43.4) | -14,151.9 (204.7) |  | -31.7 (9.6) | -8,649.2 (53.1) |
| Two T (9, 15) | -92.1 (28.1) | -15,492.8 (102.6) |  | -114.6 (45.6) | -14,169.1 (207.1) |  | -34.6 (9.7) | -8,652.1 (53.2) |
| Two T (9, 16) | -83.3 (28.0) | -15,483.9 (103.1) |  | **-56.9 (39.1)** | **-14,111.4 (197.3)** |  | -38.5 (9.8) | -8,655.9 (53.3) |
| Two T (9, 17) | -61.7 (27.5) | -15,462.4 (102.4) |  | -90.9 (41.1) | -14,145.4 (202.7) |  | -33.6 (9.8) | -8,651.0 (53.1) |

**Table S2**. Coefficient estimates from “Age + Environment” GLMMs. Coefficients in bold have 95% Bayesian credible intervals (BCI) distinct from zero. Threshold ages (T_1_, T_2_) are 7 and 15 for Breeding Date, 7 and 13 for A-egg Volume, and 6 and 13 for Clutch Size (Table S1). Unsupported quadratic effects of SSTA were removed from each model to evaluate SSTA as a linear effect. (F Age – T_1_)_+_ represents the product of (F Age – T_1_) and a logical function equal to 1 when F Age ≥ T_1_, and 0 otherwise. Conditional R^2^ includes, and marginal R^2^ excludes, variance components for Female Identity and Breeding Season.

|  | Breeding Date |  | A-egg Volume |  | Clutch Size |
| --- | --- | --- | --- | --- | --- |
| Coefficient | Mean [95% BCI] |  | Mean [95% BCI] |  | Mean [95% BCI] |
| Intercept | 2.21 [1.54, 2.86] |  | -1.44 [-1.66, -1.22] |  | -2.66 [-3.42, -1.88] |
| F Age | **-4.24 [-4.43, -4.05]** |  | **2.27 [2.06, 2.47]** |  | **6.02 [4.98, 7.06]** |
| SSTA | -0.04 [-0.33, 0.25] |  | 0.04 [-0.03, 0.11] |  | **0.30 [0.10, 0.49]** |
| SSTA^2^ | - |  | - |  | - |
| Fish Phase (Flying Fish) | 0.48 [-0.24, 1.22] |  | -0.14 [-0.34, 0.06] |  | -0.04 [-0.59, 0.56] |
| Nest Count | 0.11 [-0.24, 0.45] |  | 0.02 [-0.04, 0.07] |  | **0.27 [0.12, 0.43]** |
| FS_(_*_t_*_-1)_ | **0.40 [0.38, 0.43]** |  | 0.02 [-0.01, 0.04] |  | **0.12 [0.02, 0.21]** |
| rBD | NA |  | **-0.06 [-0.08, -0.05]** |  | -0.03 [-0.07, 0.01] |
| (F Age – T_1_)_+_ | **3.76 [3.55, 3.98]** |  | **-1.87 [-2.12, -1.64]** |  | **-5.59 [-6.71, -4.45]** |
| (F Age – T_2_)_+_ | **1.11 [0.97, 1.26]** |  | **-1.02 [-1.16, -0.88]** |  | **-1.48 [-1.89, -1.07]** |
| Group-specific effects | Var. [95% BCI] |  | Var. [95% BCI] |  | Var. [95% BCI] |
| Female ID | 0.20 [0.18, 0.21] |  | 0.60 [0.56, 0.64] |  | 0.26 [0.19, 0.33] |
| Br. Season | 0.39 [0.19, 0.84] |  | 0.03 [0.01, 0.06] |  | 0.21 [0.10, 0.46] |
| Residual | 0.40 [0.39, 0.41] |  | 0.58 [0.57, 0.59] |  |  |
| Variance explained by the model |  |  |  |  |  |
| Marginal R^2^ | 0.34 [0.25, 0.51] |  | 0.16 [0.13, 0.19] |  | 0.06 [0.04, 0.08] |
| Conditional R^2^ | 0.60 [0.59, 0.61] |  | 0.66 [0.65, 0.67] |  | 0.11 [0.10, 0.13] |

**Table S3**. Coefficient estimates from “Male Age and Identity” GLMMs. Coefficients in bold have 95% Bayesian credible intervals (BCI) distinct from zero. Threshold ages (T_1_, T_2_) are 7 and 15 for Breeding Date, 7 and 13 for A-egg Volume, and 6 and 13 for Clutch Size (Table S1). Unsupported quadratic effects of SSTA and F Age by M Age interactions were removed from models to show statistical support for lower order terms of interest. (F Age – T_1_)_+_ represents the product of (F Age – T_1_) and a logical function equal to 1 when F Age ≥ T_1_, and 0 otherwise. “x” denotes an interaction.

|  | Breeding Date |  | Clutch Size |  | A-egg Volume |  | Fledging Success |
| --- | --- | --- | --- | --- | --- | --- | --- |
| **Coefficient** | Mean [95% BCI] |  | Mean [95% BCI] |  | Mean [95% BCI] |  | Mean [95% BCI] |
| Intercept | 1.75 [0.99, 2.50] |  | -2.56 [-3.67, -1.50] |  | -1.51 [-1.82, -1.21] |  | -2.08 [-3.47, -0.62] |
| F Age | **-3.77 [-4.25, -3.30]** |  | **6.05 [4.61, 7.50]** |  | **2.29 [2.01, 2.57]** |  | **4.12 [3.11, 5.22]** |
| SSTA | 0.01 [-0.31, 0.31] |  | 0.17 [-0.07, 0.40] |  | 0.02 [-0.04, 0.09] |  | -0.26 [-0.76, 0.20] |
| SSTA^2^ | - |  | - |  | - |  | - |
| Fish Phase (Flying Fish) | 0.44 [-0.34, 1.16] |  | 0.06 [-0.70, 0.83] |  | -0.10 [-0.36, 0.15] |  | **-1.44 [-2.80, -0.08]** |
| Nest Count | 0.13 [-0.22, 0.51] |  | **0.30 [0.09, 0.56]** |  | 0.04 [-0.02, 0.10] |  | **0.51 [0.09, 1.03]** |
| FS_(_*_t_*_-1)_ | **0.44 [0.40, 0.47]** |  | 0.04 [-0.10, 0.17] |  | 0.01 [-0.03, 0.04] |  | **0.29 [0.16, 0.43]** |
| rBD | NA |  | -0.02 [-0.08, 0.04] |  | **-0.06 [-0.08, -0.04]** |  | **-0.13 [-0.19, -0.07]** |
| CS | NA |  | NA |  | NA |  | **0.67 [0.55, 0.80]** |
| rAV | NA |  | NA |  | NA |  | **0.08 [0.02, 0.14]** |
| M Age (old) | -0.10 [-0.78, 0.59] |  | -0.09 [-0.28, 0.11] |  | 0.03 [-0.03, 0.09] |  | **-0.40 [-0.59, -0.20]** |
| M Age (young) | **0.41 [0.04, 0.78]** |  | **-0.21 [-0.34, -0.08]** |  | -0.03 [-0.07, 0.01] |  | -0.10 [-0.23, 0.02] |
| (F Age – T_1_)_+_ | **3.45 [2.90, 3.98]** |  | **-5.70 [-7.28, -4.16]** |  | **-1.85 [-2.16, -1.52]** |  | **-4.33 [-5.57, -3.18]** |
| (F Age – T_2_)_+_ | **0.55 [0.23, 0.85]** |  | **-1.35 [-1.94, -0.76]** |  | **-1.04 [-1.25, -0.84]** |  | **-2.07 [-2.87, -1.31]** |
| F Age x M Age (old) | 0.11 [-0.94, 1.19] |  | - |  | - |  | - |
| F Age x M Age (young) | -0.37 [-0.91, 0.19] |  | - |  | - |  | - |
| (F Age – T_1_)_+_ x M Age (old) | -0.13 [-1.34, 1.09] |  | - |  | - |  | - |
| (F Age – T_1_)_+_ x M Age (young) | 0.23 [-0.41, 0.86] |  | - |  | - |  | - |
| (F Age – T_2_)_+_ x M Age (old) | -0.10 [-0.66, 0.51] |  | - |  | - |  | - |
| (F Age – T_2_)_+_ x M Age (young) | **0.76 [0.28, 1.24]** |  | - |  | - |  | - |
| Group-specific effects | Var. [95% BCI] |  | Var. [95% BCI] |  | Var. [95% BCI] |  | Var. [95% BCI] |
| Male ID | 0.07 [0.05, 0.09] |  | - |  | - |  | - |
| Female ID | 0.20 [0.17, 0.22] |  | 0.33 [0.21, 0.47] |  | 0.59 [0.55, 0.64] |  | 0.33 [0.20, 0.48] |
| Breeding Season | 0.41 [0.19, 0.88] |  | 0.28 [0.10, 0.65] |  | 0.02 [0.01, 0.04] |  | 1.21 [0.46, 2.73] |
| Residual | 0.36 [0.35, 0.38] |  |  |  | 0.32 [0.30, 0.33] |  |  |

**Table S4.** Coefficient estimates describing interactions between Female Age and interannual variation in the El Niño-Southern Oscillation (SSTA) affecting breeding traits in **young to middle-aged** Nazca boobies (ages 3-12). Coefficients in bold have 95% Bayesian credible intervals (BCI) distinct from zero. The threshold age dividing early-life and prime-age periods is age 9 for Breeding Probability, age 7 for Breeding Date, A-egg Volume, and Fledging Success, age 6 for Clutch Size. Models have been reduced by removing insignificant effects of SSTA^2^, as a main effect and in interactions, to show effects of SSTA. (F Age – T_1_)_+_ represents the product of (F Age – T_1_) and a logical function equal to 1 when F Age ≥ T_1_, and 0 otherwise. “x” denotes an interaction.

|  | Breeding Prob. | Breeding Date | A-egg Volume | Clutch Size | Fledging Success | FS (no earlier traits) |
| --- | --- | --- | --- | --- | --- | --- |
| Coefficients | Mean [95% BCI] | Mean [95% BCI] | Mean [95% BCI] | Mean [95% BCI] | Mean [95% BCI] | Mean [95% BCI] |
| Intercept | -4.37 [-5.27, -3.49] | 2.15 [1.54, 2.75] | -1.40 [-1.61, -1.18] | -2.80 [-3.57, -2.03] | -2.88 [-3.92, -1.77] | -3.02 [-4.06, -1.99] |
| F Age | **8.41 [7.93, 8.92]** | **-4.07 [-4.26, -3.88]** | **2.35 [2.14, 2.57]** | **6.27 [5.17, 7.30]** | **4.48 [3.65, 5.30]** | **5.44 [4.63, 6.24]** |
| SSTA | 0.59 [-0.17, 1.27] | **-0.32 [-0.62, -0.02]** | -0.03 [-0.15, 0.08] | -0.16 [-0.64, 0.33] | 0.10 [-0.59, 0.80] | 0.10 [-0.59, 0.78] |
| SSTA^2^ | -0.31 [-0.62, 0.01] | - | - | - | -0.25 [-0.55, 0.03] | -0.24 [-0.52, 0.03] |
| Fish Phase (Flying Fish) | -0.11 [-1.04, 0.78] | 0.42 [-0.29, 1.11] | **-0.25 [-0.45, -0.06]** | -0.03 [-0.58, 0.53] | -0.88 [-1.89, 0.07] | -0.85 [-1.87, 0.16] |
| Nest Count | 0.18 [-0.31, 0.64] | 0.12 [-0.23, 0.47] | 0.03 [-0.03, 0.08] | **0.27 [0.10, 0.43]** | **0.32 [0.00, 0.64]** | **0.35 [0.02, 0.69]** |
| FS_(_*_t_*_-1)_ | **-0.16 [-0.3, -0.01]** | **0.36 [0.33, 0.39]** | **0.03 [0.00, 0.06]** | **0.12 [0.02, 0.23]** | **0.36 [0.24, 0.49]** | **0.32 [0.19, 0.44]** |
| rBD | NA | NA | **-0.05 [-0.06, -0.03]** | 0.01 [-0.04, 0.06] | **-0.12 [-0.17, -0.07]** | NA |
| CS | NA | NA | NA | NA | **0.79 [0.68, 0.89]** | NA |
| rAV | NA | NA | NA | NA | **0.05 [0.00, 0.09]** | NA |
| (F Age – T_1_)_+_ | **-9.07 [-10.3, -7.89]** | **3.48 [3.22, 3.73]** | **-1.92 [-2.18, -1.67]** | **-5.82 [-7.03, -4.59]** | **-4.46 [-5.45, -3.48]** | **-5.24 [-6.22, -4.24]** |
| F Age x SSTA | **-0.47 [-1.01, 0.06]** | **0.45 [0.27, 0.61]** | 0.13 [-0.04, 0.31] | **0.94 [0.06, 1.83]** | -0.08 [-0.75, 0.58] | -0.03 [-0.66, 0.65] |
| (F Age – T_1_)_+_ x SSTA | **2.32 [0.59, 4.05]** | **-0.43 [-0.66, -0.19]** | -0.13 [-0.37, 0.10] | -1.07 [-2.19, 0.03] | 0.22 [-0.62, 1.11] | 0.12 [-0.74, 0.99] |
| F Age x SSTA^2^ | **0.66 [0.38, 0.93]** | - | - | - | - | - |
| (F Age – T_1_)_+_ x SSTA^2^ | **-2.76 [-3.65, -1.86]** | - | - | - | - | - |
| Group-specific effects | Var. [95% BCI] | Var. [95% BCI] | Var. [95% BCI] | Var. [95% BCI] | Var. [95% BCI] | Var. [95% BCI] |
| Female ID | 1.36 [1.15, 1.58] | 0.22 [0.20, 0.24] | 0.60 [0.56, 0.64] | 0.25 [0.16, 0.35] | 0.25 [0.14, 0.37] | 0.26 [0.16, 0.38] |
| Year | 0.58 [0.24, 1.36] | 0.40 [0.18, 0.84] | 0.02 [0.01, 0.05] | 0.21 [0.09, 0.44] | 0.80 [0.37, 1.72] | 0.80 [0.37, 1.69] |
| Residual |  | 0.37 [0.36, 0.39] | 0.33 [0.32 ,0.34] |  |  |  |

**Table S5.** Coefficient estimates describing interactions between Female Age and interannual variation in Nest Count (a proxy for environmental quality, Table 1) affecting breeding traits in **young to middle-aged** Nazca boobies (ages 3-12). Results are presented as in Table S4.

|  | Breeding Prob. | Breeding Date | A-egg Volume | Clutch Size | Fledging Success | FS (no earlier traits) |
| --- | --- | --- | --- | --- | --- | --- |
| Coefficients | Mean [95% BCI] | Mean [95% BCI] | Mean [95% BCI] | Mean [95% BCI] | Mean [95% BCI] | Mean [95% BCI] |
| Intercept | -4.51 [-5.30, -3.71] | 2.10 [1.45, 2.74] | -1.42 [-1.64, -1.20] | -2.76 [-3.56, -2.01] | -2.96 [-4.00, -1.87] | -3.12 [-4.18, -2.08] |
| F Age | **8.80 [8.39, 9.24]** | **-4.04 [-4.24, -3.84]** | **2.31 [2.09, 2.51]** | **6.28 [5.23, 7.36]** | **4.62 [3.78, 5.49]** | **5.64 [4.84, 6.41]** |
| SSTA | 0.33 [-0.05, 0.71] | -0.03 [-0.32, 0.24] | 0.05 [-0.01, 0.12] | **0.34 [0.15, 0.54]** | 0.09 [-0.47, 0.67] | 0.12 [-0.41, 0.65] |
| SSTA^2^ | - | - | - | - | -0.26 [-0.54, 0.01] | -0.24 [-0.52, 0.05] |
| Fish Phase (Flying Fish) | **-0.11 [-1.01, 0.78]** | 0.44 [-0.31, 1.21] | **-0.21 [-0.41, -0.01]** | -0.09 [-0.64, 0.48] | -0.87 [-1.91, 0.09] | -0.88 [-1.88, 0.16] |
| Nest Count | **0.82 [0.32, 1.28]** | 0.20 [-0.19, 0.58] | 0.00 [-0.14, 0.14] | **0.70 [0.11, 1.32]** | **0.77 [0.13, 1.41]** | **0.92 [0.32, 1.51]** |
| FS_(_*_t_*_-1)_ | **-0.15 [-0.29, -0.01]** | **0.37 [0.34, 0.40]** | **0.03 [0.00, 0.05]** | **0.13 [0.02, 0.24]** | **0.37 [0.24, 0.49]** | **0.32 [0.19, 0.45]** |
| rBD | NA | NA | **-0.05 [-0.06, -0.03]** | 0.00 [-0.05, 0.05] | **-0.12 [-0.17, -0.07]** | NA |
| CS | NA | NA | NA | NA | **0.78 [0.68, 0.88]** | NA |
| rAV | NA | NA | NA | NA | **0.05 [0.00, 0.09]** | NA |
| (F Age – T_1_)_+_ | **-11.48 [-12.48, -10.48]** | **3.45 [3.17, 3.72]** | **-1.79 [-2.07, -1.50]** | **-5.89 [-7.09, -4.69]** | **-4.64 [-5.68, -3.64]** | **-5.49 [-6.45, -4.54]** |
| F Age x Nest Count | **-1.03 [-1.41, -0.67]** | -0.07 [-0.28, 0.13] | 0.01 [-0.20, 0.21] | -0.56 [-1.62, 0.46] | -0.61 [-1.44, 0.27] | -0.80 [-1.60, 0.02] |
| (F Age – T_1_)_+_ x Nest Count | **1.50 [0.44, 2.52]** | -0.05 [-0.31, 0.23] | 0.15 [-0.11, 0.41] | 0.20 [-0.94, 1.37] | 0.41 [-0.70, 1.47] | 0.61 [-0.41, 1.62] |
| Group-specific effects | Var. [95% BCI] | Var. [95% BCI] | Var. [95% BCI] | Var. [95% BCI] | Var. [95% BCI] | Var. [95% BCI] |
| Female ID | 1.27 [1.08, 1.49] | 0.22 [0.20, 0.24] | 0.60 [0.56, 0.64] | 0.25 [0.16, 0.35] | 0.25 [0.14, 0.37] | 0.26 [0.14, 0.37] |
| Year | 0.54 [0.23, 1.19] | 0.40 [0.19, 0.84] | 0.02 [0.01, 0.05] | 0.21 [0.09, 0.43] | 0.80 [0.38, 1.61] | 0.81 [0.39, 1.60] |
| Residual |  | 0.38 [0.37 ,0.39] | 0.33 [0.32 ,0.34] |  |  |  |

**Table S6.** Coefficient estimates describing interactions between Female Age and interannual variation in the El Niño-Southern Oscillation (SSTA) affecting breeding traits in **middle-aged to old** Nazca boobies (ages 11-20). The threshold age (T_2_) dividing prime-age and late-life periods is at 16 years for Breeding Probability, 15 years for Breeding Date and Fledging Success, and 13 years for Clutch Size and A-egg Volume. Results presented as in Table S4.

|  | Breeding Prob. | Breeding Date | A-egg Volume | Clutch Size | Fledging Success | FS (no earlier traits) |
| --- | --- | --- | --- | --- | --- | --- |
| Coefficients | Mean [95% BCI] | Mean [95% BCI] | Mean [95% BCI] | Mean [95% BCI] | Mean [95% BCI] | Mean [95% BCI] |
| Intercept | 2.79 [1.49, 4.14] | 0.51 [-0.18, 1.21] | 0.18 [-0.20, 0.55] | 1.75 [0.41, 3.09] | 0.49 [-0.62, 1.64] | 0.98 [-0.14, 2.08] |
| F Age | -0.15 [-0.97, 0.66] | **-0.32 [-0.59, -0.07]** | -0.09 [-0.40, 0.23] | -0.46 [-1.47, 0.58] | **-0.94 [-1.50, -0.34]** | **-0.93 [-1.49, -0.33]** |
| SSTA | 0.46 [-0.82, 1.69] | **1.02 [0.32, 1.67]** | 0.16 [-0.15, 0.47] | 1.01 [-0.19, 2.22] | 0.13 [-0.81, 1.05] | 0.16 [-0.74, 1.05] |
| SSTA^2^ | **-** | -0.36 [-0.75, 0.01] | - | - | - | - |
| FS_(_*_t_*_-1)_ | 0.15 [-0.09, 0.38] | **0.50 [0.46, 0.54]** | -0.02 [-0.06, 0.03] | -0.01 [-0.15, 0.14] | **0.39 [0.21, 0.56]** | **0.26 [0.08, 0.44]** |
| rBD | NA | NA | **-0.07 [-0.09, -0.04]** | -0.02 [-0.09, 0.05] | **-0.21 [-0.28, -0.13]** | NA |
| CS | NA | NA | NA | NA | **0.63 [0.47, 0.79]** | NA |
| rAV | NA | NA | NA | NA | **0.07 [0.00, 0.15]** | NA |
| (F Age – T_1_)_+_ | -0.90 [-2.90, 1.10] | **0.54 [0.08, 1.02]** | **-0.46 [-0.86, -0.07]** | -0.32 [-1.60, 1.04] | -0.97 [-2.13, 0.17] | -1.05 [-2.25, 0.09] |
| F Age x SSTA | -0.25 [-1.07, 0.55] | **-0.27 [-0.53, -0.02]** | -0.15 [-0.40, 0.09] | -0.59 [-1.50, 0.32] | -0.21 [-0.66, 0.25] | -0.20 [-0.63, 0.24] |
| F Age x SSTA^2^ | - | 0.07 [-0.06, 0.20] | - | - | - | - |
| (F Age – T_1_)_+_ x SSTA | 0.82 [-1.09, 2.70] | **0.77 [0.25, 1.27]** | 0.09 [-0.22, 0.40] | 0.45 [-0.66, 1.60] | 0.03 [-0.88, 0.92] | -0.04 [-0.94, 0.85] |
| (F Age – T_1_)_+_ x SSTA^2^ | - | -0.20 [-0.45, 0.05] | - | - | - | - |
| Group-specific effects | Var. [95% BCI] | Var. [95% BCI] | Var. [95% BCI] | Var. [95% BCI] | Var. [95% BCI] | Var. [95% BCI] |
| Female ID | 2.29 [1.72, 2.98] | 0.24 [0.21, 0.27] | 0.66 [0.61, 0.72] | 0.35 [0.17, 0.54] | 0.29 [0.10, 0.53] | 0.28 [0.08, 0.50] |
| Year | 1.07 [0.27, 3.40] | 0.36 [0.09, 1.17] | 0.03 [0.01, 0.10] | 0.43 [0.11, 1.34] | 1.38 [0.44, 3.89] | 1.42 [0.43, 4.00] |
| Residual |  | 0.38 [0.37, 0.40] | 0.31 [0.30, 0.33] |  |  |  |

**Table S7.** Coefficient estimates describing interactions between Female Age and interannual variation in Nest Count (a proxy for environmental quality, Table 1) affecting breeding traits in **middle-aged to old** Nazca boobies (ages 11-20). The threshold age (T_2_) dividing prime-age and late-life periods is at 16 years for Breeding Probability, 15 years for Breeding Date and Fledging Success and 13 years for Clutch Size and A-egg Volume. Results presented as in Table S4.

|  | Breeding Prob. | Breeding Date | A-egg Volume | Clutch Size | Fledging Success | FS (no earlier traits) |
| --- | --- | --- | --- | --- | --- | --- |
| Coefficients | Mean [95% BCI] | Mean [95% BCI] | Mean [95% BCI] | Mean [95% BCI] | Mean [95% BCI] | Mean [95% BCI] |
| Intercept | 2.73 [1.46, 3.97] | 0.14 [-0.42, 0.73] | 0.06 [-0.31, 0.43] | 1.51 [0.08, 2.91] | 0.54 [-0.61, 1.68] | 1.00 [-0.12, 2.19] |
| F Age | -0.18 [-0.96, 0.60] | **-0.31 [-0.48, -0.13]** | 0.02 [-0.29, 0.31] | -0.30 [-1.36, 0.74] | **-0.97 [-1.57, -0.40]** | **-0.95 [-1.49, -0.38]** |
| Nest Count | -0.33 [-1.54, 0.80] | -0.35 [-0.87, 0.20] | 0.17 [-0.23, 0.57] | -0.52 [-1.77, 0.62] | -0.74 [-1.81, 0.35] | -0.70 [-1.73, 0.29] |
| FS_(_*_t_*_-1)_ | 0.15 [-0.09, 0.39] | **0.50 [0.45, 0.54]** | -0.01 [-0.06, 0.03] | -0.01 [-0.17, 0.15] | **0.38 [0.19, 0.56]** | **0.26 [0.09, 0.44]** |
| rBD | NA | NA | **-0.07 [-0.09, -0.04]** | -0.02 [-0.10, 0.05] | **-0.21 [-0.28, -0.13]** | NA |
| CS | NA | NA | NA | NA | **0.62 [0.46, 0.77]** | NA |
| rAV | NA | NA | NA | NA | **0.07 [0.00, 0.14]** | NA |
| (F Age – T_1_)_+_ | -0.73 [-2.68, 1.22] | **0.43 [0.10, 0.77]** | **-0.57 [-0.93, -0.20]** | -0.50 [-1.83, 0.82] | -0.98 [-2.15, 0.19] | **-1.09 [-2.25, 0.00]** |
| F Age x Nest Count | 0.44 [-0.23, 1.13] | 0.09 [-0.07, 0.26] | -0.08 [-0.41, 0.22] | 0.47 [-0.42, 1.44] | **0.62 [0.08, 1.16]** | **0.59 [0.06, 1.12]** |
| (F Age – T_1_)_+_ x Nest Count | -0.83 [-2.58, 0.93] | 0.03 [-0.33, 0.35] | -0.03 [-0.44, 0.40] | -0.52 [-1.76, 0.67] | -0.94 [-2.04, 0.20] | -0.92 [-2.04, 0.23] |
| Group-specific effects | Var. [95% BCI] | Var. [95% BCI] | Var. [95% BCI] | Var. [95% BCI] | Var. [95% BCI] | Var. [95% BCI] |
| Female ID | 2.28 [1.67, 3.00] | 0.24 [0.21, 0.27] | 0.66 [0.61, 0.72] | 0.34 [0.18, 0.51] | 0.29 [0.08, 0.53] | 0.27 [0.07, 0.50] |
| Year | 1.09 [0.28, 3.47] | 0.54 [0.16, 1.62] | 0.02 [0.01, 0.08] | 0.56 [0.16, 1.66] | 1.51 [0.44, 4.44] | 1.48 [0.45, 4.58] |
| Residual |  | 0.38 [0.37, 0.40] | 0.31 [0.30, 0.33] |  |  |  |
