## Appendix S9 for "Breeding responses to environmental variation are age- and trait-dependent in female Nazca boobies"

### APPENDIX S9 – Results: late breeding in seasons 2013-2016

For the Age + Environment GLMMs explaining variation in Breeding Date, posterior predictive checking of model fit revealed that random Breeding Season effects for 2013, 2014, and 2015 (and, to a lesser extent, 2016) were larger than expected under the assumption that all Breeding Season effects were drawn from the same Gaussian distribution (Fig. S1), leading us to consider *a posteriori* the existence of an unrecognized environmental effect applicable to these four seasons (when breeding was unusually late, Fig. S2). Interannual variation in the Pacific Decadal Oscillation (PDO) covaries with Fish Phase and influences adult survival in Nazca boobies (Champagnon et al. 2019). Notably, the PDO index shifted from negative to positive anomalies in January 2014 (during the 2013 Breeding Season) and maintained that status until a return to neutral conditions in late 2017/ early 2018. We downloaded monthly values of the PDO (<http://jisao.washington.edu/pdo/PDO.latest>) for the years 1992-2018 and calculated a 12-month average (October-September, following the Nazca booby breeding schedule). We then regressed breeding season PDO on SSTA and used the resulting residuals as a predictor of Breeding Date (residual PDO is uncorrelated with SSTA) in place of Fish Phase in Age + Environment GLMMs. After updating Age + Environment models, the distribution of random Breeding Season effects fell closer to the models’ assumptions (not shown) and effects of female age, SSTA, and Nest Count on Breeding Date were unchanged (Table S1), confirming that SSTA and Nest Count do not influence average Breeding Date.

**Table S1**. Coefficient estimates from the best-supported Age + Environment GLMM for Breeding Date after replacing Fish Phase with a predictor capturing interannual variation in the Pacific Decadal Oscillation (PDO). First and second threshold ages are 7 and 15, respectively. Coefficients in bold have 95% Bayesian credible intervals (BCI) distinct from zero. Unsupported quadratic effects of SSTA were removed from each model to evaluate SSTA as a linear effect. (F Age – T_1_)_+_ represents the product of (F Age – T_1_) and a logical function equal to 1 when F Age ≥ T_1_, and 0 otherwise. Conditional R^2^ includes, and marginal R^2^ excludes, variance components for Female Identity and Breeding Season.

|  | Breeding Date (with PDO) |
| --- | --- |
| Coefficient | Mean [95% BCI] |
| Intercept | 2.57 [2.26, 2.87] |
| F Age | **-4.24 [-4.43, -4.05]** |
| SSTA | 0.06 [-0.22, 0.32] |
| SSTA^2^ | - |
| PDO | 0.40 [-0.01, 0.83] |
| Nest Count | 0.26 [-0.11, 0.65] |
| FS_(_*_t_*_-1)_ | **0.40 [0.37, 0.43]** |
| (F Age – T_1_)_+_ | **3.77 [3.55, 3.98]** |
| (F Age – T_2_)_+_ | **1.11 [0.96, 1.26]** |
| Group-specific effects | Var. [95% BCI] |
| Female ID | 0.20 [0.18, 0.21] |
| Breeding Season | 0.34 [0.16, 0.71] |
| Residual | 0.40 [0.39, 0.41] |
| Variance explained by model |  |
| Marginal R^2^ | 0.38 [0.26, 0.57] |
| Conditional R^2^ | 0.60 [0.59, 0.61] |

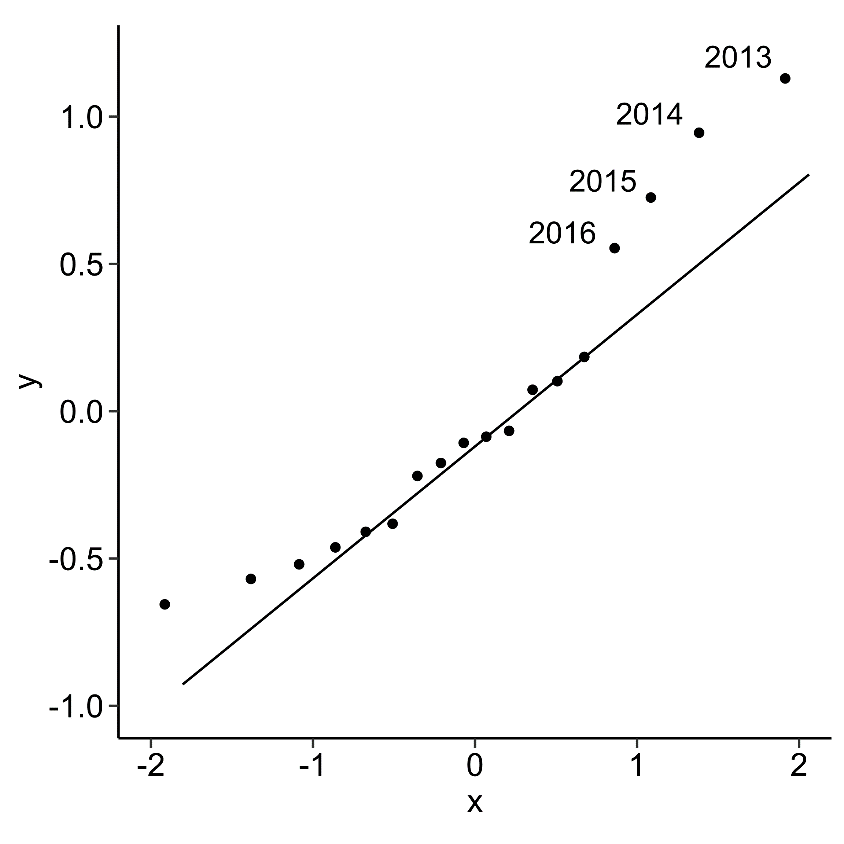

**Figure S1**. Quantile-Quantile plot showing the distribution of random breeding season effects estimated in the best-supported Age + Environment GLMM explaining variation in Breeding Date (Table 2) plotted against theoretical quantiles drawn from a standard normal distribution.

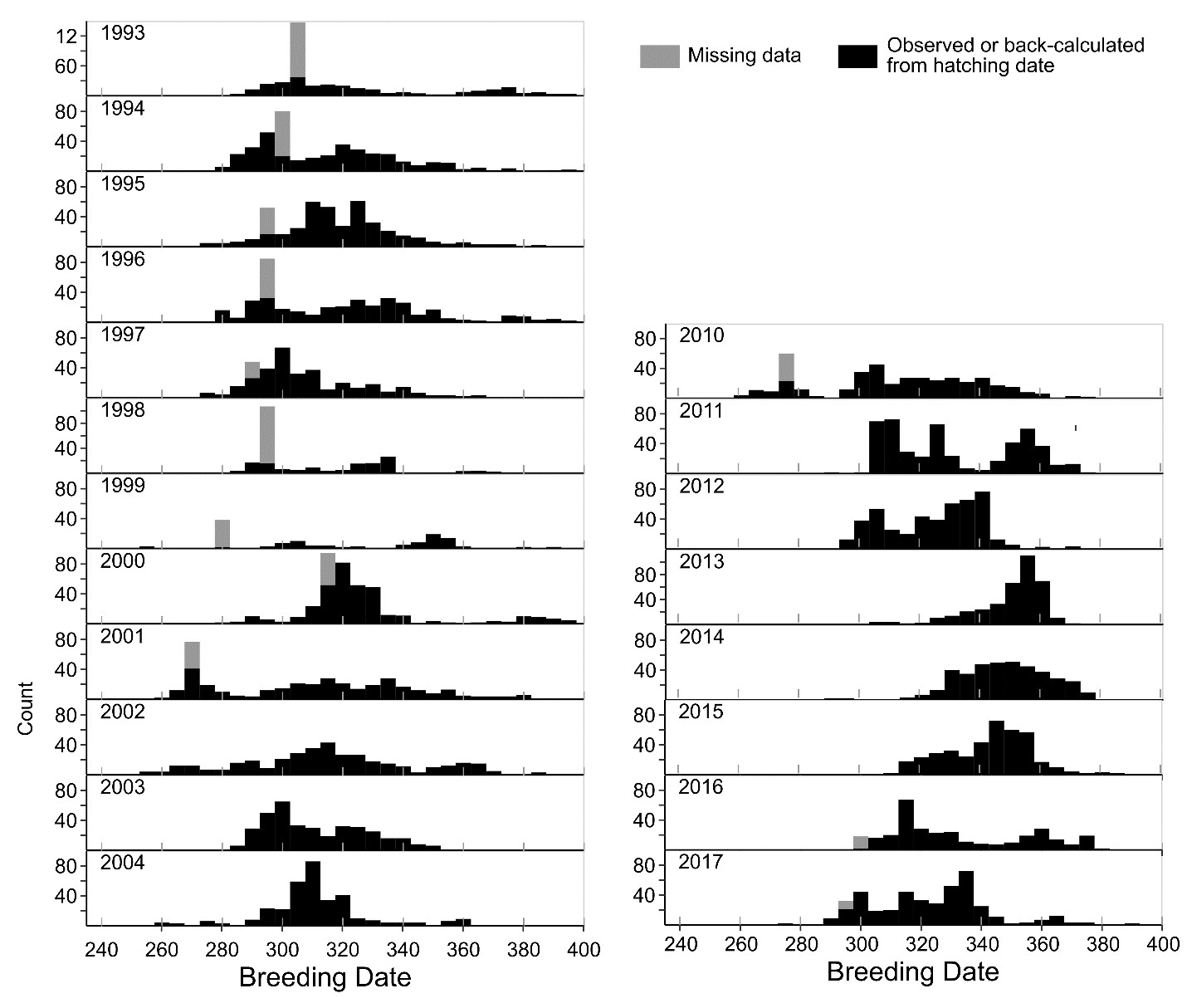
 **Figure S2**. Breeding Dates by Breeding Season. Data come from the Study Area, a comprehensively monitored colony subsection. Breeding Dates are in extended Julian dates. Breeding Dates directly observed, or back-calculated from hatching dates, are in black. Breeding Dates of clutches laid before colony monitors arrived and failing to hatch were estimated as the median Breeding Date of samples laid before monitors arrived but hatching (shown in grey; Appendix S1).
