## Appendix S10 for "Breeding responses to environmental variation are age- and trait-dependent in female Nazca boobies"

### APPENDIX S10 – Accounting for heterogeneous error variances when modelling Breeding Date

Posterior predictive checks on models fit to Breeding Date suggested that the raw data are more variable in some years than in others (Fig. S1), violating the assumption that within-group errors are independent and identically normally distributed (Pinheiro and Bates 2000). Additional models were fit to Breeding Date during the Age + Environment analysis stage to evaluate the sensitivity of our inferences to the models’ violations of assumptions (Gelman et al. 2013). We performed model checking early in the study, before an additional two Breeding Seasons (2016 and 2017) were added to the data, explaining why the results shown below exclude those two years.

Table S1 compares coefficient values estimated when including unequal error variance by year and when failing to do so for the Age + Environment candidate model including linear + quadratic age effects. Both models were fit in R (v. 3.4.3, R Core Team 2017) using package MCMCglmm (v. 2.25, Hadfield 2010) because rstanarm (v. 2.13.1, Goodrich et al. 2016) does not accommodate unequal error variances. MCMCglmm uses Gibbs sampling to estimate the posterior distribution. We ran a single chain per model for 2,000,000 iterations (after a burn-in period of 50,000) with a thinning interval of 1,000. We thus retained 2,000 posterior samples per model, which were used to calculate marginal posterior means and credible intervals for coefficients. Priors were identical to those used in our original methods (Appendix S6) except for Female Identity and Breeding Season random effects, which received parameter-expanded priors (following Hadfield 2011). From Table S1, the heteroscedastic model fit the data better than the homoscedastic model (ΔDIC = 2,580), but coefficient estimates on fixed effects and variance components had similar magnitudes and directions. Results were similar for a comparison of heteroscadastic vs. homoscedastic models from the Male Age and Identity analyses and the Age by Environment Interaction analyses (not shown), and lead to the same conclusion: the magnitude and direction of effects of interest were affected little by a failure to model year-specific error variance, increasing confidence in our original GAMMs and GLMMs (reported in the main text).

*References*

Gelman, A., J. B. Carlin, H. S. Stern, D. B. Dunson, A. Vehtari, and D. B. Rubin. 2013. *Bayesian Data Analysis*, Third Edit. CRC Press, Boca Raton, FL.

Goodrich, B., J. Gabry, I. Ali & S. Brilleman. 2016. rstanarm: Bayesian applied regression modeling via Stan. R package version 2.13.1. <http://mc-stan.org/>.

Hadfield, J. D. 2010. [MCMC methods for multi-response generalized linear mixed models: the MCMCglmm R package](http://www.jstatsoft.org/v33/i02/paper). *Journal of Statistical Software,* **33**, 1-22.

R Core Team. 2017. R: A language and environment for statistical computing. R Foundation for Statistical Computing, Vienna, Austria.

**Table S1**. Coefficient estimates describing additive effects of female age and environment on Breeding Date (Stage 1 analyses) comparing the results of ignoring (homoscedastic model), versus accounting for, heteroscedasticity. Coefficients in bold have 95% Bayesian credible intervals (BCI) distinct from zero.

|  | Homoscedastic model |  |  | Heteroscedastic model |
| --- | --- | --- | --- | --- |
| Coefficient | Mean [95% BCI] |  | Coefficient | Mean [95% BCI] |
| Intercept | -1.46 [-2.89, -0.08] |  | Intercept | -1.35 [-2.68, -0.03] |
| **F Age** | **-0.42 [-0.43, -0.40]** |  | **F Age** | **-0.35 [-0.37, -0.34]** |
| **F Age^2^** | **0.17 [0.16, 0.18]** |  | **F Age^2^** | **0.14 [0.13, 0.14]** |
| SSTA | 0.08 [-0.35, 0.56] |  | SSTA | 0.07 [-0.39, 0.54] |
| SSTA^2^ | -0.05 [-0.27, 0.19] |  | SSTA^2^ | -0.04 [-0.26, 0.20] |
| Fish Phase (Flying Fish) | 0.53 [-0.30, 1.30] |  | Fish Phase (Flying Fish) | 0.50 [-0.31, 1.26] |
| Nest Count | 0.07 [-0.16, 0.31] |  | Nest Count | 0.08 [-0.14, 0.34] |
| **FS_(_*_t_* _- 1)_** | **0.30 [0.27, 0.32]** |  | **FS_(_*_t_* _- 1)_** | **0.25 [0.23, 0.28]** |
| Group-specific effects | Var. [95% BCI] |  | Group-specific effects | Var. [95% BCI] |
| Female ID | 0.16 [0.15, 0.18] |  | Female ID | 0.13[0.12, 0.14] |
| Br. Season | 0.44 [0.16, 0.84] |  | Br. Season | 0.43[0.16, 0.83] |
| Residual | 0.40 [0.39, 0.41] |  | Residual 1993 | 0.62 [0.47, 0.77] |
|  |  |  | Residual 1994 | 0.35 [0.27, 0.44] |
|  |  |  | Residual 1995 | 0.27 [0.21, 0.35] |
|  |  |  | Residual 1996 | 0.83 [0.68, 1.01] |
|  |  |  | Residual 1997 | 0.31 [0.24, 0.37] |
|  |  |  | Residual 1998 | 0.53 [0.4, 0.66] |
|  |  |  | Residual 1999 | 1.65 [1.16, 2.18] |
|  |  |  | Residual 2000 | 0.41 [0.34, 0.48] |
|  |  |  | Residual 2001 | 1.29 [1.1, 1.49] |
|  |  |  | Residual 2002 | 0.98 [0.86, 1.12] |
|  |  |  | Residual 2003 | 0.38 [0.33, 0.44] |
|  |  |  | Residual 2004 | 0.44 [0.38, 0.5] |
|  |  |  | Residual 2010 | 0.85 [0.78, 0.91] |
|  |  |  | Residual 2011 | 0.5 [0.46, 0.53] |
|  |  |  | Residual 2012 | 0.23 [0.21, 0.25] |
|  |  |  | Residual 2013 | 0.12 [0.11, 0.13] |
|  |  |  | Residual 2014 | 0.18 [0.16, 0.2] |
|  |  |  | Residual 2015 | 0.18 [0.16, 0.19] |
| DIC | 25,350.82 |  | DIC | 22,770.85 |

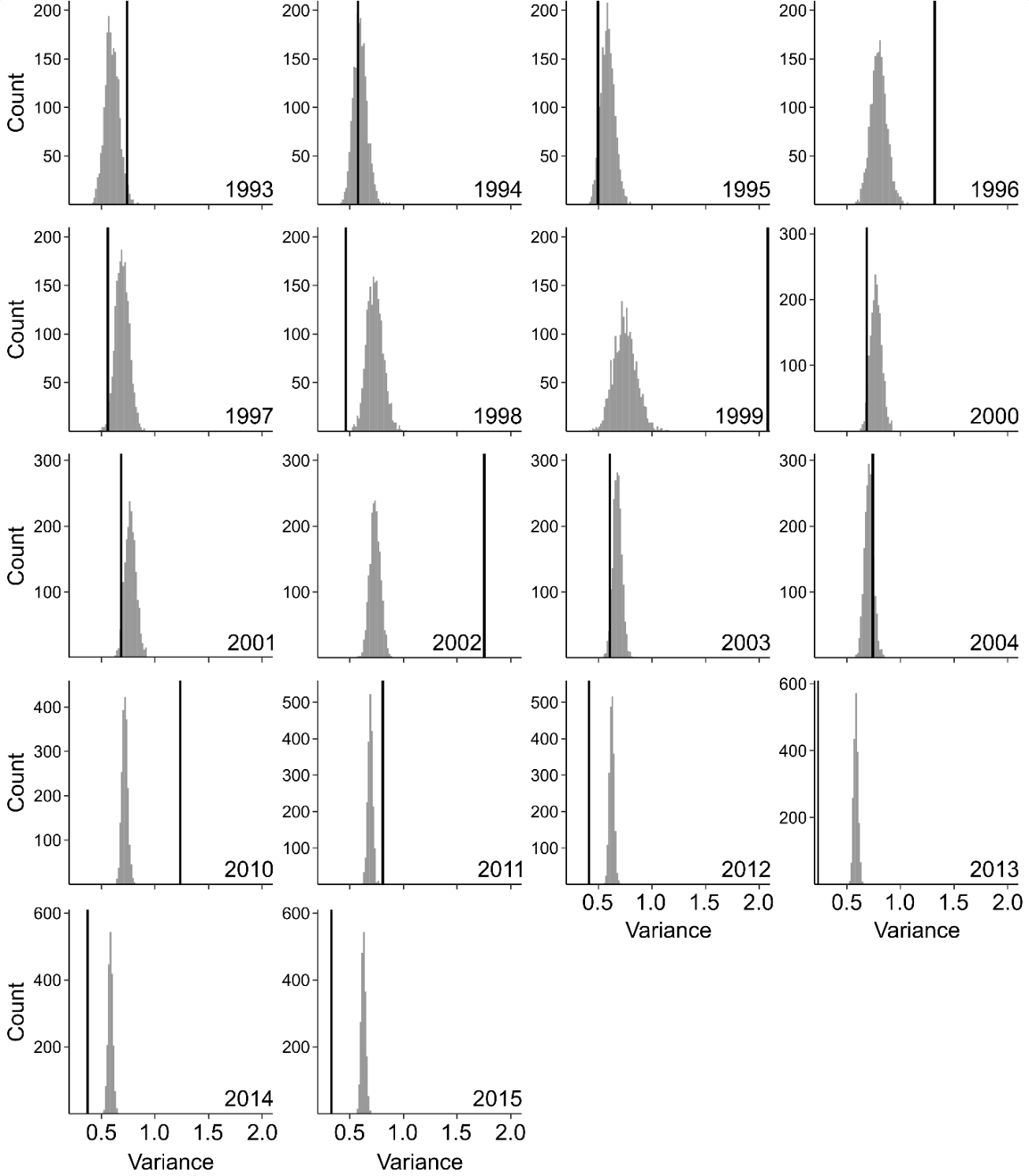

**Figure S1**. Posterior predicted variance in Breeding Date by breeding season (grey histograms), from the Age + Environment GAMM (main text Table 2), fits the original data poorly for many years (black lines show year-specific Breeding Date variances). Histograms show the distribution from 1,000 posterior predictive samples.
